# Mechanistic Insights into Specific G Protein Interactions with Adenosine Receptors Revealed by Accelerated Molecular Simulations

**DOI:** 10.1101/541250

**Authors:** Jinan Wang, Yinglong Miao

**Affiliations:** Center for Computational Biology and Department of Molecular Biosciences, University of Kansas, Lawrence, KS 66047, USA

## Abstract

Coupling between G-protein-coupled receptors (GPCRs) and the G proteins is a key step in cellular signaling. Despite extensive experimental and computational studies, the mechanism of specific GPCR-G protein coupling remains poorly understood. This has greatly hindered effective drug design of GPCRs that are primary targets of ~1/3 of currently marketed drugs. Here, we have employed all-atom molecular simulations using a robust Gaussian accelerated molecular dynamics (GaMD) method to decipher the mechanism of the GPCR-G protein interactions. Adenosine receptors (ARs) were used as model systems based on very recently determined cryo-EM structures of the A_1_AR and A_2A_AR coupled with the G_i_ and G_s_ proteins, respectively. Changing the G_i_ protein to the G_s_ led to increased fluctuations in the A_1_AR and agonist adenosine (ADO), while agonist 5’-N-ethylcarboxamidoadenosine (NECA) binding in the A_2A_AR could be still stabilized upon changing the G_s_ protein to the G_i_. Free energy calculations identified one stable low-energy conformation for each of the ADO-A_1_AR-G_i_ and NECA-A_2A_AR-G_s_ complexes as in the cryo-EM structures, similarly for the NECA-A_2A_AR-G_i_ complex. In contrast, the ADO agonist and G_s_ protein sampled multiple conformations in the ADO-A_1_AR-G_s_ system. GaMD simulations thus indicated that the ADO-bound A_1_AR preferred to couple with the G_i_ protein to the G_s_, while the A_2A_AR could couple with both the G_s_ and G_i_ proteins, being highly consistent with experimental findings of the ARs. More importantly, detailed analysis of the atomic simulations showed that the specific AR-G protein coupling resulted from remarkably complementary residue interactions at the protein interface, involving mainly the receptor transmembrane 6 helix and the Gα α5 helix and α4-β6 loop. In summary, the GaMD simulations have provided unprecedented insights into the dynamic mechanism of specific GPCR-G protein interactions at an atomistic level, which is expected to facilitate future drug design efforts of the GPCRs.

## Introduction

G-protein-coupled receptors (GPCRs) are key cellular signaling proteins and represent primary targets of ~1/3 of currently marketed drugs.^1^ Particularly, four subtypes (A_1_, A_2A_, A_2B_, and A_3_) of GPCRs mediate the effects of adenosine, an endogenous nucleoside modulator that plays a critical role in cytoprotective function. Adenosine receptors (ARs) have emerged as important therapeutic targets for treating many human diseases such as cardiac ischemia, neuropathic pain and cancer.^2^ During function, the A_1_AR and A_3_AR preferentially couple to the G_i/o_ proteins, while the A_2A_AR and A_2B_AR preferentially couple to the G_s_ proteins. Nevertheless, increasing evidence suggests that GPCRs including the ARs can couple to multiple G proteins.^3–6^

Few complex structures of GPCRs coupled with the G protein or its mimic have been determined using X-ray crystallography or cryo-EM so far.^7^ ARs are the sole subfamily of GPCRs that have structures in complex with different G proteins, *i.e.*, the adenosine (ADO)-bound A_1_AR coupled with the G_i_ protein^8^ and the 5’-N-ethylcarboxamidoadenosine (NECA)-bound A_2A_AR coupled with an engineered G_s_ protein.^9^ Both structures were obtained via cutting-edge cryo-EM and published very recently in 2018. The GPCR-G protein complex structures provide valuable information about active conformations of the GPCRs and G proteins. However, they are rather static images. The dynamic mechanism of specific GPCR–G protein interactions remains unclear. Experimental techniques including mutagenesis, nuclear magnetic resonance, hydrogen-deuterium exchange mass spectrometry, double electron-electron resonance spectroscopy and structural biology have been utilized to investigate GPCR–G protein interactions.^10–13^ While the C-terminal α_5_ helix in the G_α_ subunit has been suggested as the primary driver for specific receptor recognition, the G_α_ α_N_ helix and receptor intracellular loop (ICL) 2 and transmembrane (TM) 6 helix further contribute to the GPCR–G protein coupling specificity. In addition, dynamic regions in the complex and agonist binding can be crucial for the coupling through allosteric modulation.^11,12^

A bioinformatics approach has been applied to determine a selectivity barcode (patterns of amino acids) of GPCR–G protein coupling.^14^ While universally conserved residues in the barcode allow GPCRs to bind and activate G protein in a similar manner, different receptors recognize the unique positions of the G-protein barcode through distinct residues. Molecular dynamics (MD) simulations have identified several important regions for coupling of the G protein with activated GPCRs, including the receptor TM6 and G_α_ α5 helices.^15–17^ MD simulations have shown that conformational dynamics of the GPCR-G protein complex depends on the bound ligands.^18,19^ Moreover, MD simulations have suggested that the binding of the active GPCR is necessary for nucleotide release from the G protein.^20–23^ However, due to limited timescales, conventional MD (cMD) simulations often suffer from insufficient sampling, precluding proper free energy calculations to characterize GPCR-G protein interactions quantitatively.

To overcome the limitations of cMD, enhanced sampling methods have been applied to investigate GPCR–G protein interactions. Umbrella sampling has been used to calculate free energy profiles of the TM6 outward movement during receptor coupling to the G proteins.^17^ Metadynamics simulations have been performed to investigate the dynamic effects of different GPCR ligands and intracellular binding partners^24^ and examine differences of GPCRs coupled by the G protein versus its mimetic nanobody.^25^ Nevertheless, these enhanced simulation methods require predefined collective variables and may apply constrains on the conformational space of the proteins. In this regard, a novel and robust Gaussian accelerated MD (GaMD) method has been developed to allow for unconstrained enhanced sampling and free energy calculations of large biomolecules^26–28^. GaMD has been applied to successfully simulate protein folding^26,27^, protein-ligand binding and unbinding^26,27,29^, GPCR activation^29^, large-scale conformational transitions of the CRISPR-Cas9 gene-editing system^30^, T cell receptor signaling protein^31^ and human dystonia related protein^32^, and so on. Notably, GaMD has been recently applied to capture spontaneous binding of the G-protein mimetic nanobody to a muscarinic GPCR.^33^

In this study, we have employed all-atom enhanced sampling simulations using the robust GaMD method on the latest cryo-EM structures of the ADO-A_1_AR-G_i_ and NECA-A_2A_AR-G_s_ protein complexes, as well as complexes with the G proteins switched (**Table 1** and **Figure S1**). A computational model was prepared for the receptor-G protein complexes in explicit lipids and solvent (**Figure S2**). The GaMD simulations allowed us to characterize structural flexibility and low-energy conformations of the AR-G protein complexes, which provided important insights into the mechanism of specific GPCR–G protein interactions.

**Table 1.** Summary of GaMD simulations performed on the agonist-bound AR-G protein complexes. ^a^ The AR:NPxxY-G:α5 distance is the center-of-mass (COM) distance between the receptor NPxxY motif and the last 5 residues of the G_α_ α5 helix. b The R3.50-E6.30 distance is the distance between the Cα atoms of conserved residues Arg^3.50^ and Glu^6.30^ in the receptors. ^c^ Theα5 orientation angle is the angle between COMs of the receptor orthosteric pocket, the last 5 and first 5 residues of the G_α_ α5 helix, illustrated in **Figure S8**. ^d^ The increase of G_α_-G_β_ distance is the increase in the distance between the COMs of G_α_ (excluding the N-terminal helix) and G_β_ (excluding the C-terminal of β sheet) subunits compared to the cryo-EM structure.

| System | ADO-A<sub>1</sub>AR-G<sub>i</sub> | NECA-A<sub>2A</sub>AR-G<sub>s</sub> | NECA-A<sub>2A</sub>AR-G<sub>i</sub> | ADO-A<sub>1</sub>AR-G<sub>s</sub> |
| --- | --- | --- | --- | --- |
| Dimension (Å<sup>3</sup>) | 112x124x150 | 110x124x148 | 110x124x148 | 108x124x153 |
| *N*<sub>atoms</sub> | 180,394 | 175,140 | 176,028 | 176,940 |
| Simulation Length (ns) | 300 x 3 | 300 x 3 | 300 x 3 | 300 x 3 |
| Boost Potential (kcal/mol) | 21.43 ± 6.50 | 21.12 ± 6.49 | 20.96 ± 6.41 | 21.88 ± 6.58 |
| ***Cryo-EM experimental structures*** |  |  |  |  |
| AR:NPxxY-G:α5 Distance<sup>a</sup> (Å) | 12.7 | 13.6 | - | - |
| R3.50-E6.30 Distance<sup>b</sup> (Å) | 16.3 | 18.4 | - | - |
| α5 Orientation Angle <sup>c</sup> (°) | 143.9 | 123.5 | - | - |
| ***GaMD simulation low-energy conformations*** |  |  |  |  |
| AR:NPxxY-G:α5 Distance (Å) | 12.2 | 13.2 | 11.2 | 11.8, 13.5 |
| R3.50-E6.30 Distance (Å) | 17.4 | 18.2 | 17.6 | 17.5, 22.5 |
| α5 Orientation Angle (°) | 135.2 | 127.5 | 140.2 | 123.0, 130.0 |
| Increase of G<sub>α</sub>-G<sub>β</sub> Distance<sup>d</sup> (Å) | -1.3 | 1.5 | -0.8 | 1.2, 3.2 |

## Results

### Variations of structural flexibility in different adenosine receptor-G protein complexes

In GaMD simulations of the ADO-A_1_AR-G_i_ and NECA-A_2A_AR-G_s_ complexes, the receptors underwent small fluctuations except the extracellular loop 2 (ECL2) and TM6 intracellular end (**Figure S3**). Overall, the G proteins exhibited higher flexibility than the receptors, especially in the α5 helix, α4-β6 loop and α4-β5 loop in the G_α_ subunit and terminal regions of the G_βγ_ subunits. Both the A_1_AR and A_2A_AR showed flexibility change upon switching of the G proteins. For the A_1_AR, changing the G_i_ protein to the G_s_ led to increased fluctuations in the ADO agonist and the receptor ECL2, TM6 intracellular end and helix 8 (H8) (**Figure 1A**). These motifs were suggested to be important in previous studies for activation of the A_1_AR and receptor coupling with the G protein.^8,34^ For the A_2A_AR, changing the G_s_ protein to the G_i_, however, appeared to stabilize the receptor with slightly decreased fluctuations in the latter, other than the ECL2 and intracellular loop 2 (ICL2) regions (**Figure 1B**).

**Figure 1.**
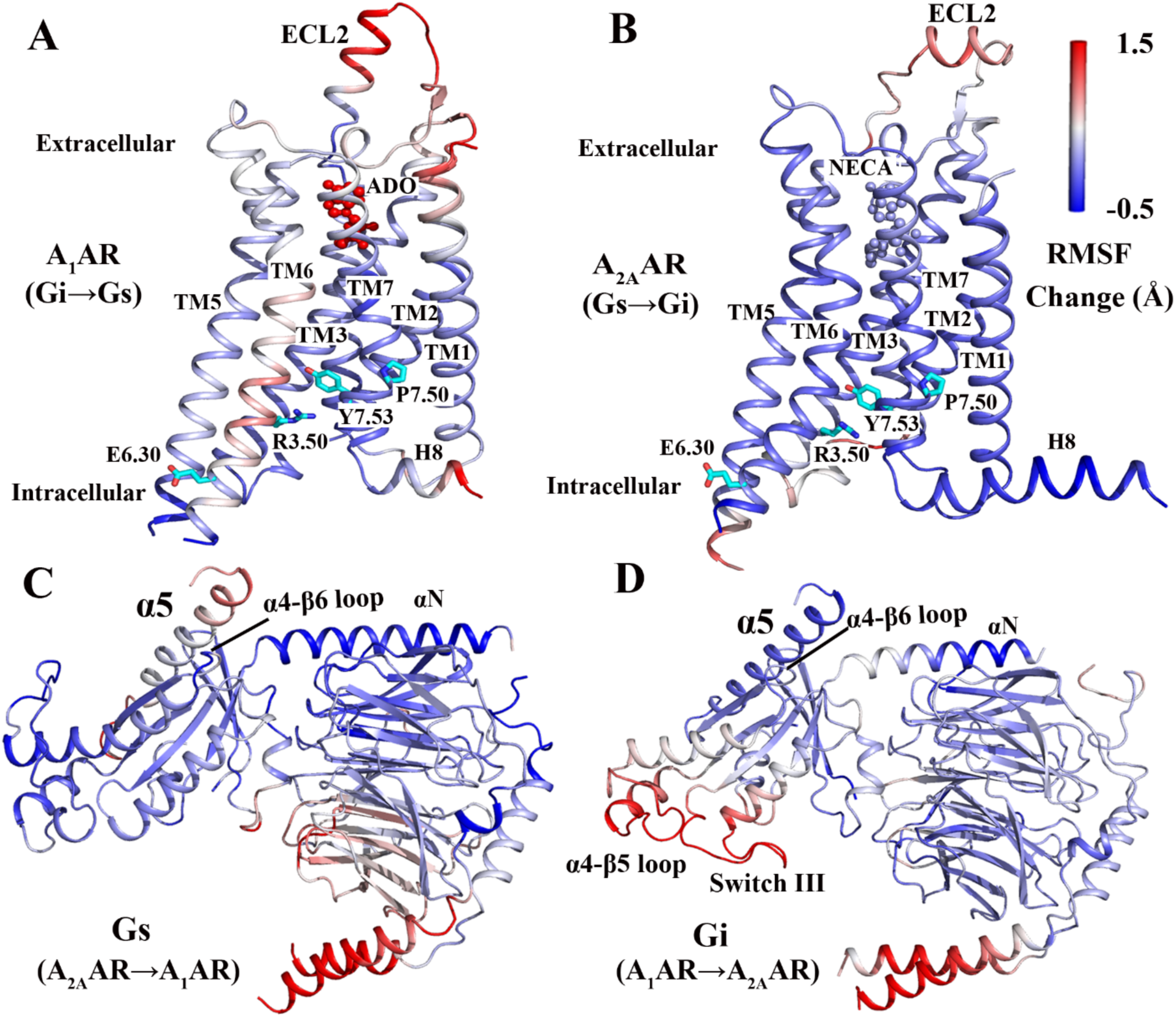
Comparison of structural flexibility of active adenosine receptor-G protein complexes obtained from GaMD simulations: **(A)** Change in the root-mean-square fluctuations (RMSFs) of the A_1_AR when the G_i_ protein was changed to the G_s_ protein. **(B)** Change in the RMSFs of the A_2A_AR when the G_s_ protein was changed to the G_i_ protein. **(C)** Change in the RMSFs of the G_s_ protein when the receptor was changed from the A_2A_AR to the A_1_AR. **(D)** Change in the RMSFs of the G_i_ protein when the receptor was changed from the A_1_AR to the A_2A_AR. A color scale of −0.5 Å (blue) to 1.5 Å (red) is used.

Next, we examined flexibility change of the G proteins upon coupling to the different receptors. In the G_s_ protein, the C-terminus of the G_α_ α5 helix exhibited higher fluctuations when the A_2A_AR was changed to the A_1_AR (**Figure 1C**). In the G_i_ protein, while the G_α_ α5 helix became stabilized with lower fluctuations, the α4-β5 loop and switch III exhibited higher flexibility when the A_1_AR was changed to the A_2A_AR (**Figure 1D**). These regions were shown earlier to play a key role in activation and receptor recognition of the G protein.^21,35^

### Distinct binding modes of the G proteins and agonists in adenosine receptors

Free energy profiles were calculated from GaMD simulations to identify low-energy conformations of the GPCR-G protein complexes. RMSD of the agonist relative to the cryo-EM structures and the distance between the receptor NPxxY motif in the TM7 intracellular end and the C-terminus of the G_α_ α5 helix were first used as the reaction coordinates. In the ADO-A_1_AR-G_i_ and NECA-A_2A_AR-G_s_ protein complexes, both the G proteins and agonists maintained their cryo-EM conformations (**Figures 2A and 2B**, **Table 1**). In the NECA-A_2A_AR-G_i_ complex, the NECA agonist maintained the cryo-EM conformation as in the NECA-A_2A_AR-G_s_ complex, but the G_i_ protein sampled a different state with the receptor:NPxxY-G_α_ α5 distance decreased to ~11.2 Å (**Figure 2C**). The G_αi_ α5 helix moved ~2 Å towards the TM7 NPxxY motif of the A_2A_AR relative to the G_αs_ α5 helix in the NECA-A_2A_AR-G_s_ structure (**Table 1**). Nevertheless, the NECA-A_2A_AR-G_i_ complex adopted a stable low-energy conformation in the free energy profile (**Figure 2C**).

In the ADO-A_1_AR-G_s_ system, the ADO agonist sampled two low-energy conformational states, denoted “L1” and “L2”, for which agonist RMSD relative to the cryo-EM conformation was ~3.0 Å and ~7.5 Å, respectively (**Figure 2D**). The “L1” conformation of ADO was similar to the cryo-EM structure with slight sliding of the purine ring by ~2 Å at the orthosteric site (**Figure 3A**). In the “L2” conformation, ADO formed interactions with residues Tyr^1.35^ and Tyr^7.36^ in the “sub-pocket 2” of the A_1_AR described earlier^36^ (**Figure S4**). Residue superscripts denote Ballesteros and Weinstein (BW) numbering of GPCRs.^37^ The G_s_ protein sampled two low-energy conformational states, which were similar to cryo-EM conformations of the G_i_ protein in the ADO-A_1_AR-G_i_ complex and the G_s_ protein in the NECA-A_2A_AR-G_s_ complex. The receptor:NPxxY-G_α_:α5 distance was ~11.8 Å and ~13.5 Å in the G_i_- and G_s_-bound A_1_AR, respectively (**Figure 2D** and **Table 1**).

### High flexibility and conformational changes of ECL2 in the A_1_AR

In the A_1_AR, different conformational states of the ECL2 helix were identified from GaMD simulations, including the “open”, “semi-open” and “closed” (**Figure 3B**). In comparison, the ECL2 in the A_2A_AR sampled only one low-energy conformational state (**Figure S5**). In the ADO-A_1_AR-G_i_ complex, the ECL2 sampled a distinct “semi-open” conformation with ~5.5 Å RMSD in the helix region compared with the starting “open” cryo-EM conformation, for which the ECL2 helix tilted towards the receptor TM bundle by ~4 Å (**Figure 3B**). An energy barrier of ~1.1 kcal/mol was found between the “open” and “semi-open” states of ECL2 in the ADO-A_1_AR-G_i_ complex (**Figure S5A**). In the ADO-A_1_AR-G_s_ system, the receptor ECL2 transitioned between the “open” and “closed” states with ~1.2 kcal/mol energy barrier (**Figure 4A**). In the “closed” state, the ECL2 helix tilted towards the receptor TM bundle by ~9 Å relative to the “open” cryo-EM conformation (**Figure 3B**). Meanwhile, the TM2 extracellular domain could move outwards by ~10 Å (**Figure S6B**). Such movement was observed in binding of a covalent antagonist DU172 to the A_1_AR that involved an induced fit mechanism.^36,38^ During conformational transition from the “open” state to the “closed” in ECL2 of the A_1_AR, residue Trp156^ECL2^ switched its hydrogen bonding partner from Gly163^ECL2^ to Val166^ECL2^ (**Figure S6**). This finding was consistent with previous mutagenesis experiments, suggesting that residues Trp156^ECL2^ and Val166^ECL2^ were important in the activation and allosteric modulation of the A_1_AR.^34,39^

### Comparatively weak coupling between the A_1_AR and G_s_ protein

Overall, the ADO-A_1_AR-G_i_, NECA-A_2A_AR-G_s_ and NECA-A_2A_AR-G_i_ complexes appeared to be stable during the GaMD simulations (**Table 1**). Each of them sampled only one low-energy conformation in the free energy profiles (**Figures 2**, **S7**, **S9** and **S10**). In comparison, coupling of the G_s_ protein to the ADO-bound A_1_AR was significantly weaker. The ADO-A_1_AR-G_s_ system deviated from the simulation starting structure, visiting multiple low-energy conformational states. In particular, the TM6 intracellular end of the A_1_AR sampled two distinct low-energy states, referred to as “Active” and “Over-active”, for which the Arg^3.50^-Glu^6.30^ distance was ~17.5 Å and ~22.5 Å, respectively (**Figure 4B**). The “Active” state exhibited ~0.8 kcal/mol lower free energy than the “Over-active” state. The receptor TM6 intracellular end moved ~5 Å away from the TM bundle in the “Over-active” state compared with the “Active” state (**Figures 3C** and **4B**). When the A_1_AR visited the “Active” and “Over-active” states, the G_α_ α5 helix adopted an orientation angle of ~123° and ~130° (**Figure 4C**) and the distance between the G_α_ and G_β_ subunits increased by ~1.2 Å and ~3.2 Å, respectively (**Figure 4D**).

The above results suggested that the G_s_ protein could not stabilize the ADO-bound A_1_AR. On the other hand, the G_αs_ and G_βs_ subunits tended to dissociate from each other when the G_s_ protein coupled to the A_1_AR. In contrast, the G_αi_ and G_βi_ subunits formed closer interaction when the G_i_ protein coupled to the ADO-bound A_1_AR, for which the G_αi_-G_βi_ distance decreased by ~1.3 Å in the energy minimum conformation of the ADO-A_1_AR-G_i_ complex (**Figure S10A**). Therefore, the A_1_AR induced closer interaction of the G_α_ and G_β_ subunits in the G_i_ protein, but dissociation of the G_αs_ and G_βs_ subunits from each other (**Table 1**). In summary, coupling of ADO-bound A_1_AR to the G_s_ protein was weaker than to the G_i_ protein.

**Figure 2.**
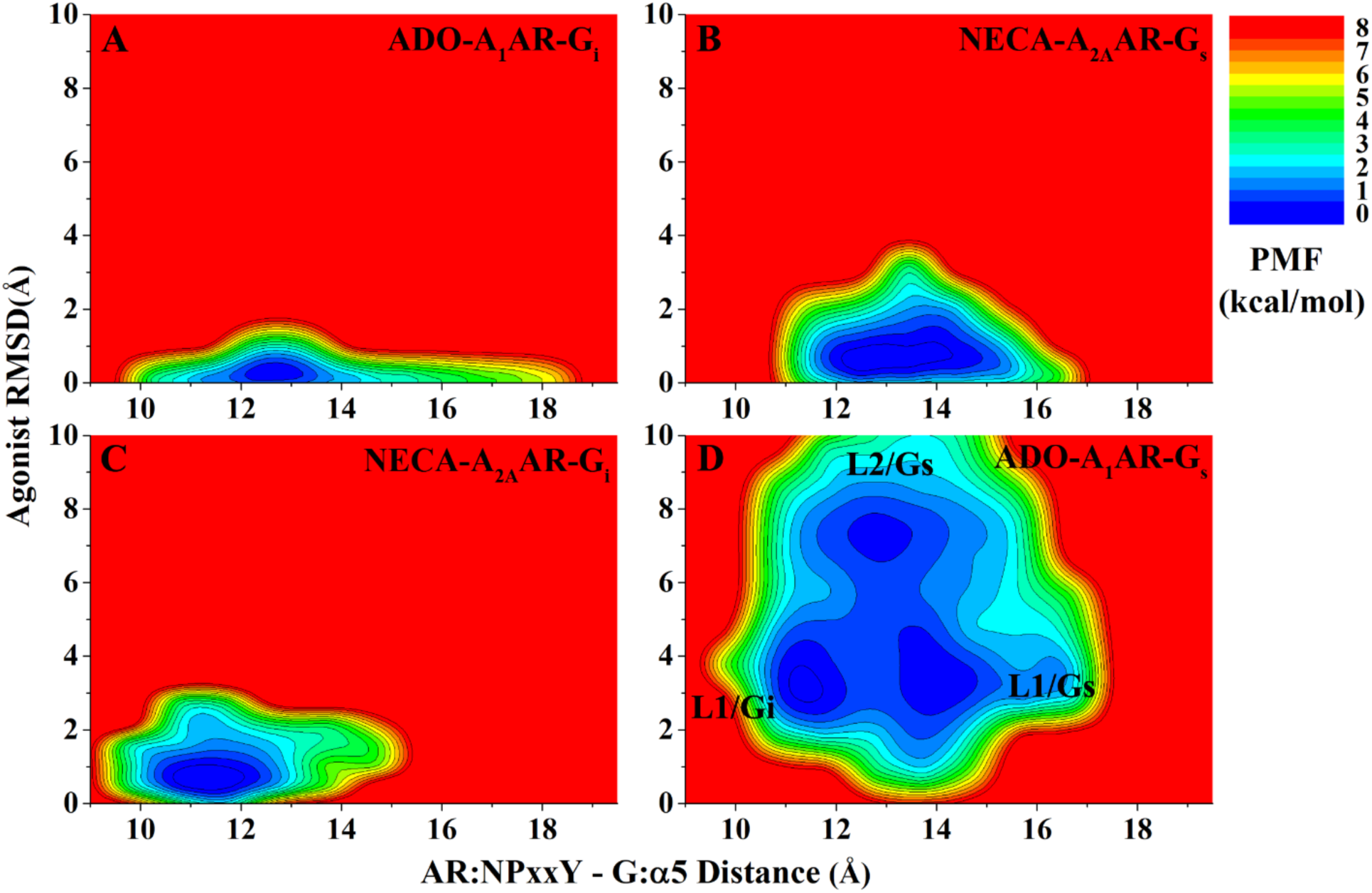
2D potential of mean force (PMF) profiles of the **(A)** ADO-A_1_AR-G_i_, **(B)** NECA-A_2A_AR-G_s_, **(C)** NECA-A_2A_AR-G_i_ and **(D)** ADO-A_1_AR-G_s_ complex systems regarding the agonist RMSD relative to the cryo-EM conformation and AR:NPxxY-G:α5 distance.

**Figure 3.**
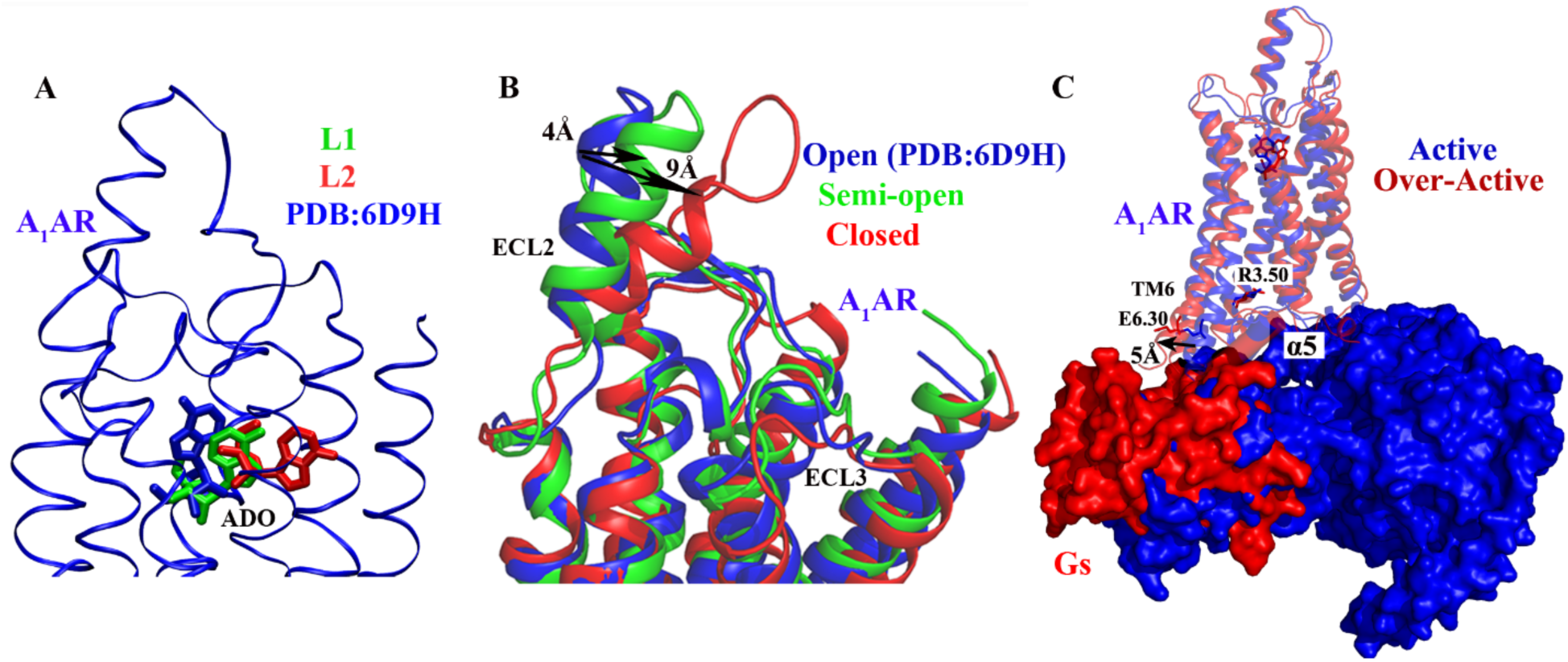
Distinct active conformations of the A_1_AR induced by binding of the G_s_ protein: **(A)** Representative conformations of two low-energy binding poses of ADO (L1 and L2 in green and red, respectively) in the A_1_AR-G_s_complex. The cryo-EM structure of the A_1_AR-G_i_ complex (PDB: 6D9H, blue) is shown for comparison. **(B)** Representative conformations of open (PDB: 6D9H, blue), semi-open (green) and closed (red) states of ECL2 in the A_1_AR. **(C)** Representative conformations of the A_1_AR in the “Active” (blue) and “Over-active” (red) states in the A_1_AR-G_s_ system.

**Figure 4.**
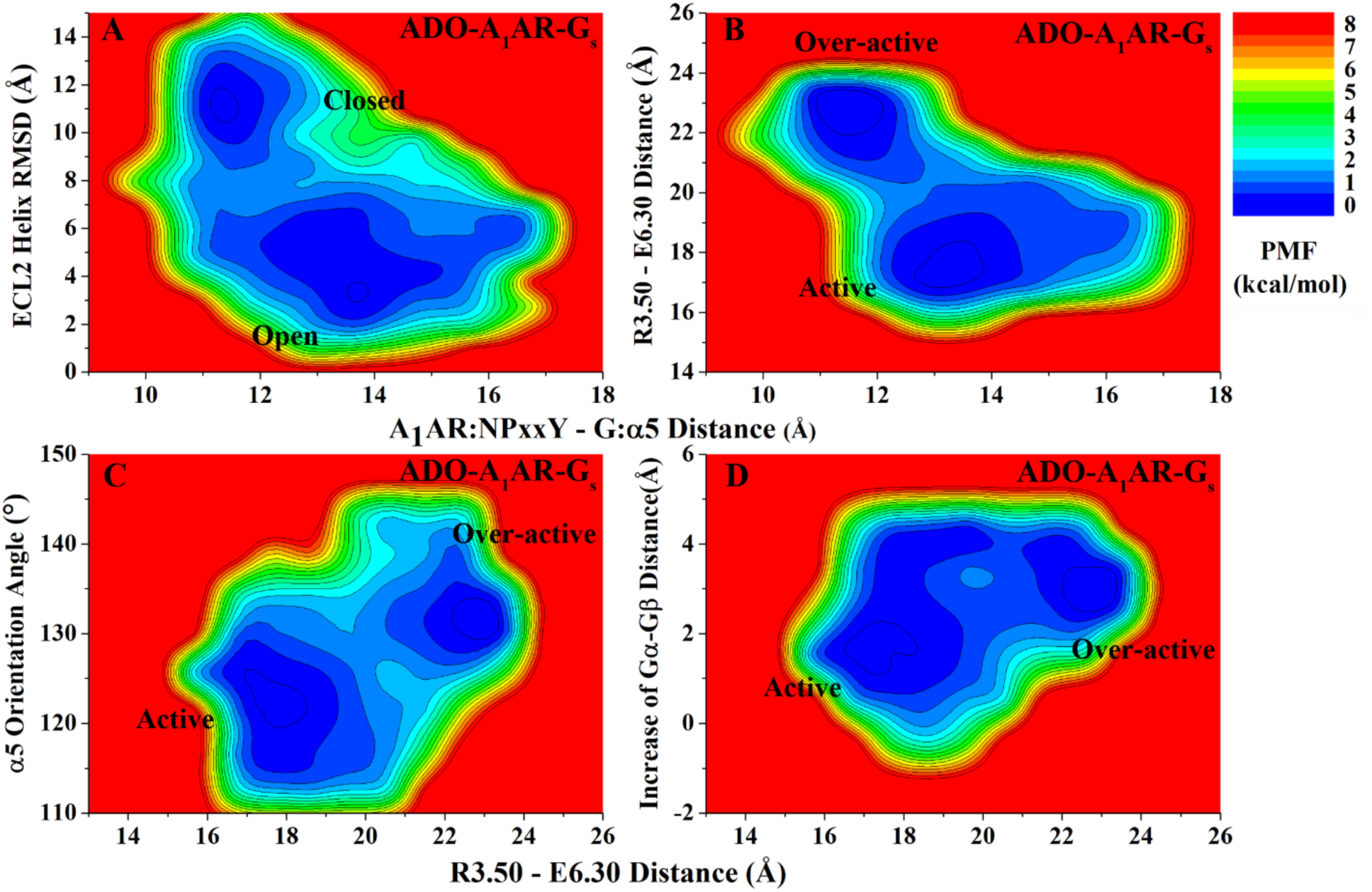
Distinct low-energy conformational states of the ADO-A_1_AR-G_s_ system were identified from GaMD simulations. **(A)** 2D PMF of RMSD of the helix region in ECL2 relative to the cryo-EM structure and the A_1_AR: NPxxY-G: α5 distance. **(B)** 2D PMF of the R3.50-E6.30 and A_1_AR: NPxxY-G: α5 distances. **(C)** 2D PMF of the orientation angle of the G_α_ α5 helix and the R3.50-E6.30 distance. **(D)** 2D PMF of the increase of the G_α_-G_β_ distance and the R3.50-E6.30 distance.

### Mechanism of specific adenosine receptor-G protein interactions

Sequence alignments of the ARs and G proteins (**Figure S11**) and detailed comparison of residue interactions at the protein interface (**Figure 5**) enabled us to identify the origin of specific AR-G protein interactions. In the A_1_AR-G_i_ and A_2A_AR-G_s_ complexes, analysis of low-energy conformations identified from the GaMD simulations highlighted specific residue interactions **(Figures 5A** and **5B**) that were similar to those obtained previously by comparing the A_1_AR-G_i_ and β2AR-G_s_ experimental structures.^8^ The last five residues of the G_α_ α5 helix formed significantly stronger receptor interactions in the A_1_AR-G_i_ complex than in the A_2A_AR-G_s_ complex. Residue Asp351 or GH5.22 with common Gα numbering (CGN) 40 in the G_i_ protein formed salt-bridge interactions with Arg^3.53^ and Lys^8.49^ in the A_1_AR. In the G_s_ protein, residue Glu382 (GH5.24) formed similar salt-bridge interactions with Arg^7.56^ and Arg^8.51^ in the A_2A_AR. However, these five residues formed closer van der Waals interactions with the TM3, TM5, TM6 and H8 of the A_1_AR. Second, residues GH5.8 – GH5.21 of the G_α_ α5 helix formed more interactions with the receptor ICL2 and TM5 helix in the A_2A_AR-G_s_ complex instead than in the A_1_AR-G_i_ complex. In addition, the α4–β6 loop of the G protein formed distinct interactions with the receptor TM5 helix in the A_2A_AR-G_s_ complex compared with the A_1_AR-G_i_ complex. Residues His347 (Gh4s6.13) and Tyr348 (Gh4s6.20) of the G_s_ protein formed non-polar interactions with Gln^5.71^ and Met^5.72^ in the A_2A_AR (**Figure 5B**). In contrast, residue Asp316 (Gh4s6.9) of the G_i_ protein formed polar interactions with residues Lys^6.25^ and Tyr^6.26^ in the A_1_AR (**Figure 5A**).

In comparison, the A_2A_AR-G_i_ complex appeared to form more residue interactions at the protein interface (**Figure 5C**) than both the A_1_AR-G_i_ and A_2A_AR-G_s_ complexes. Notably, the last five residues of G_αi_ α5 helix formed extensive polar and non-polar interactions with the TM3, TM6, TM7, ICL2 and H8 of the A_2A_AR. Residue Phe355 (GH5.26) formed close interactions with Lys^6.29^, His^6.32^ and Ser^6.36^ of the A_2A_AR. Residues GH5.8-GH5.21 of the G_αi_ α5 helix formed additional interactions with H8 of the A_2A_AR, apart from those with the receptor TM5 and ICL2 as observed in the A_1_AR-G_i_ and A_2A_AR-G_s_ complexes. Furthermore, the β2-β3 loop of the G_i_ protein formed new interactions with the ICL2 of the A_2A_AR. Residue Asp194 (Gs2s3.2) formed a salt-bridge with Arg111^ICL2^ in the A_2A_AR. These interactions greatly contributed to strong coupling of the A_2A_AR and G_i_ protein, which showed stable low-energy conformations in the free energy profiles (e.g., **Figure 2**) and small fluctuations (**Figure 1**).

**Figure 5.**
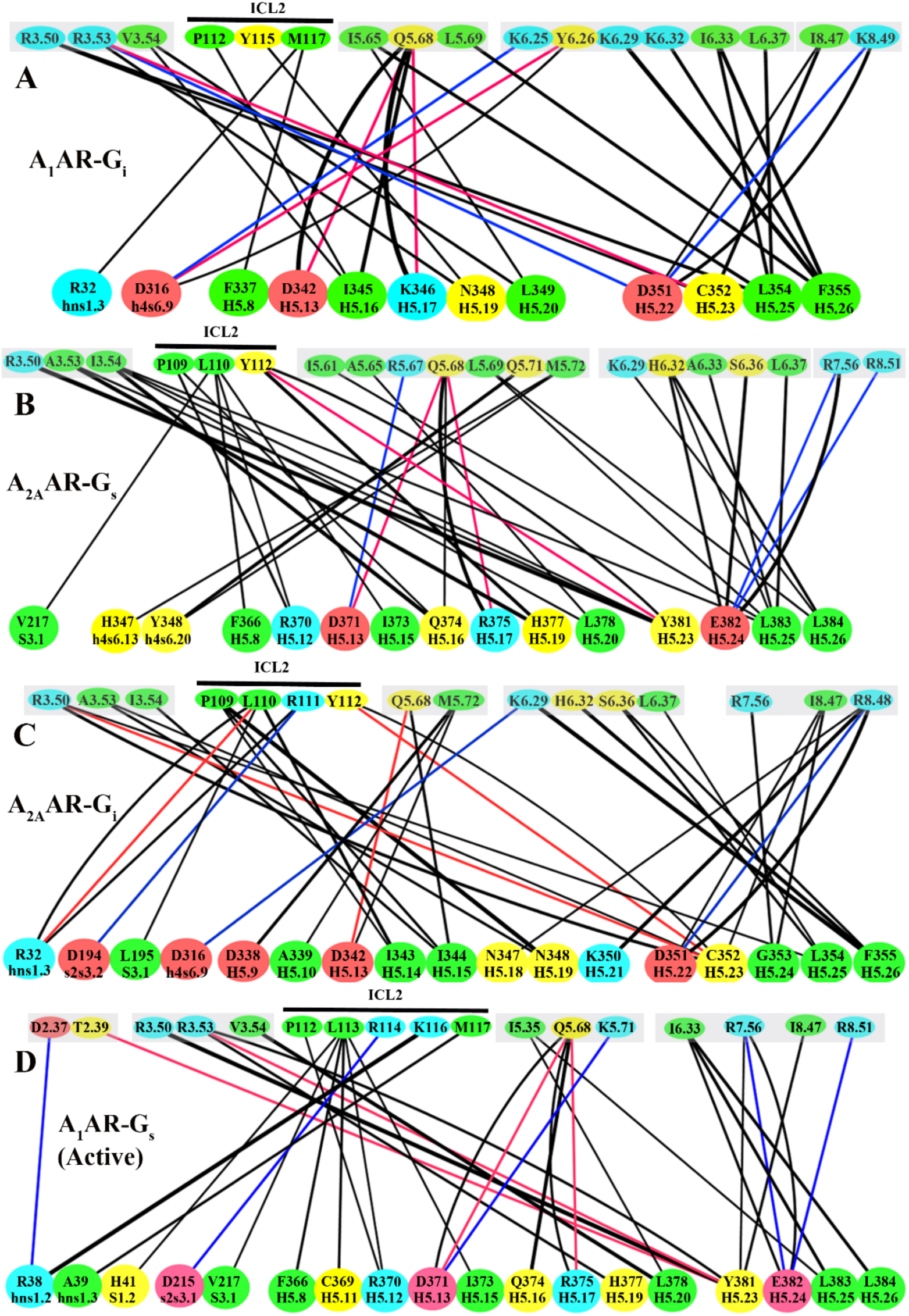
Complementary residue interactions at the protein interface in the (**A**) ADO-A_1_AR-G_i_, (**B**) NECA-A_2A_AR-G_s_, (**C**) NECA-A_2A_AR-G_i_ and (**D)** ADO-A_1_AR-G_s_ (active state) systems. Receptor and G-protein residues are numbered according to the BW and CGN schemes, respectively. Hydrogen bond, van der Waals and salt-bridge interactions are colored in red, black and blue, respectively. The line thickness is proportional to the number of residue interactions. Hydrophobic, polar, acidic and basic residues are colored in green, yellow, red and blue, respectively.

When the G_s_ protein coupled to the A_1_AR, residue interactions at the protein interface were decreased overall (**Figure 5D**). In the “Active” state, the TM6 helix of the A_1_AR formed significantly fewer interactions with the G_s_ protein (**Figure 5D**) than with the G_i_ protein (**Figure 5A**). The αN-β1, β2 sheet and β2-β3 loop of the G_s_ protein involving residues R38 (Ghns1.2), A39 (Ghns1.3), H41 (GS1.2), D215 (Gs2s3.1) and V217 (GS3.1) formed new interactions with the TM2 helix and ICL2 of the A_1_AR (**Figure 5D**). However, both clusters of residues in the G_α_ α5 helix (GH5.8-GH5.21 and GH5.22-GH5.26) greatly reduced receptor interactions in the A_1_AR-G_s_ system compared with the A_1_AR-G_i_ and A_2A_AR-G_s_ complexes. Similar results were observed in the “Over-active” conformation of the A_1_AR-G_s_ system (**Figure S12**). Therefore, reduced residue interactions were found at the protein interface between the A_1_AR and G_s_ protein, leading to their weaker coupling compared with the other three AR-G protein complexes (**Figure 5**).

In summary, the ADO-bound A_1_AR preferred to bind the G_i_ protein to the G_s_, while the A_2A_AR could bind both the G_s_ and G_i_ proteins (**Figure 6**). For the A_1_AR, when the G_i_ protein was changed to the G_s_, the receptor ECL2 and TM6 intracellular end underwent higher fluctuations and sampled multiple conformational states, similarly for the agonist and G protein. The G_s_ protein could not stabilize ADO binding in the A_1_AR, and vice versa. Coupling of the A_1_AR to the G_s_ protein was significantly weaker than to the G_i_ protein. The G_αs_ and G_βs_ subunits tended to dissociate from each other (**Figure 6A**). In contrast, both the G_s_ and G_i_ proteins could stabilize agonist NECA binding in the A_2A_AR. The G_i_ protein became more compact with decreased distance between the G_α_ and G_β_ subunits, contributing to strong coupling of the G_i_ protein to the NECA-bound A_2A_AR (**Figure 6B**).

**Figure 6.**
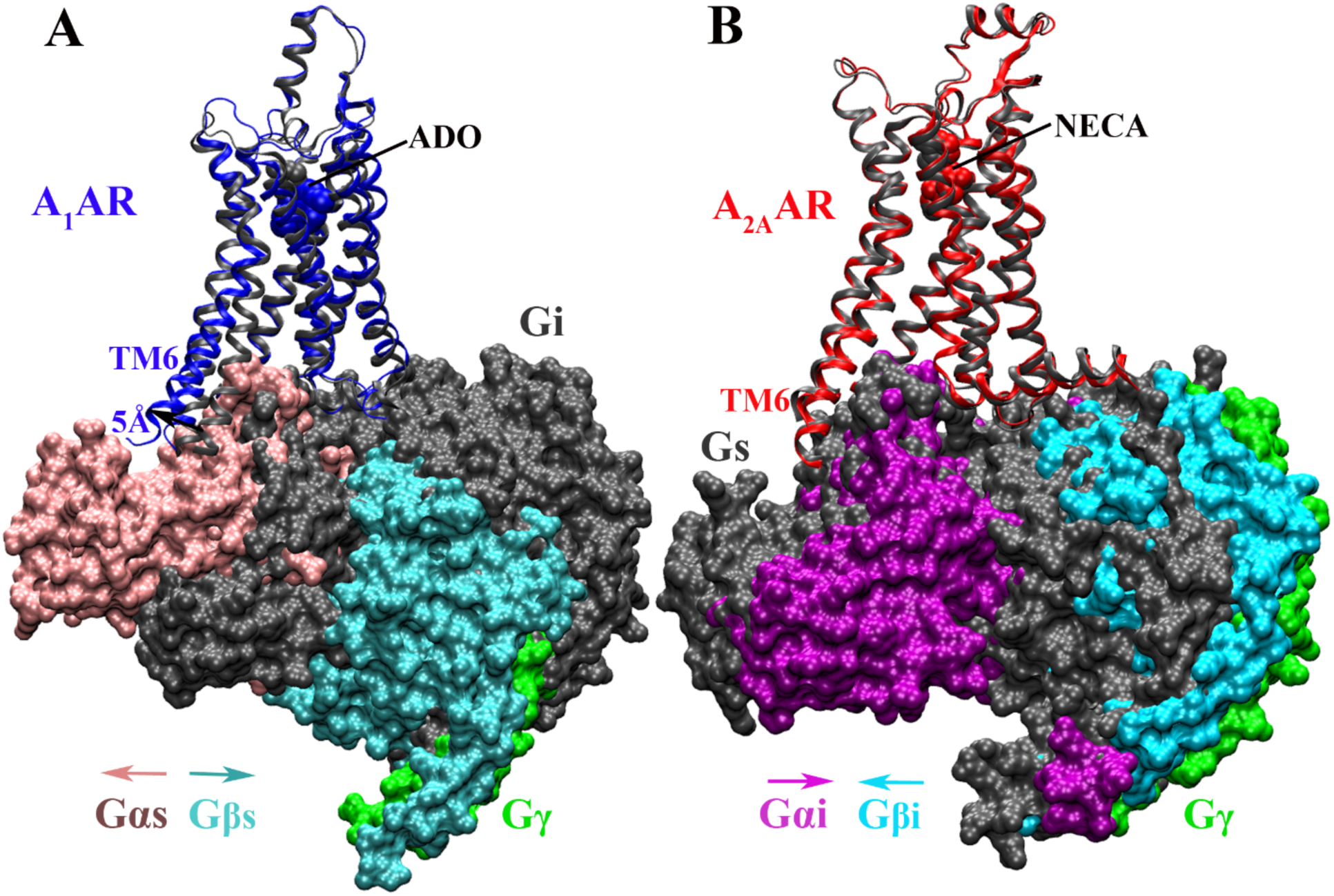
Summary of specific AR-G protein interactions: **(A)** the ADO-bound A_1_AR prefers to bind the G_i_ protein to the G_s_. The latter could not stabilize agonist ADO binding in the A_1_AR and tended to dissociate from the receptor. **(B)** The A_2A_AR could bind both the G_s_ and G_i_ proteins, which adopted distinct conformations in the complexes.

## Discussions

ARs are so far the only subfamily of GPCRs that have available experimental structures in complex with different G proteins. The AR-G protein complex structures were obtained via cutting-edge cryo-EM and published very recently in 2018. Here, all-atom GaMD enhanced simulations have unprecedentedly allowed us to characterize the structural flexibility and low-energy conformations of the different AR-G protein complexes. Further detailed analysis of the AR-G protein complex simulations has highlighted a network of remarkable complementary residue interactions at the protein interface, which are important for specific G protein coupling to the A_1_AR and A_2A_AR.

The ADO-bound A_1_AR preferred to bind the G_i_ protein to the G_s_. This was consistent with previous experimental studies that the A_1_AR coupled to different G proteins with the following rank order of “preference”: G_i_>G_s_>G_q_, and the G protein coupling depends on the bound agonist potency._4,5_ Particularly, the NECA-bound A_1_AR was shown to preferentially couple to the G_i_ protein compared with the G_s_ protein.^4^ Considering similar binding affinities of ADO and NECA in the A_1_AR,^41,42^ the ADO-bound A_1_AR likely prefers to couple to the G_i_ protein as well. When the A_1_AR coupled with the G_s_ protein, the ADO agonist exhibited high fluctuations and sampled two different binding poses (“L1” and “L2”). In the “L2” binding pose, ADO formed interactions with residues Tyr^1.35^ and Tyr^7.36^ in the sub-pocket 2 of the A_1_AR as described earlier.^36^ This was similar to the 5UIG X-ray structure of the A_2A_AR,^43^ in which the 8D1 antagonist interacted with the same residues of the A_2A_AR (**Figure S4**). The ECL2 of the A_1_AR was highly flexible (**Figures 1** and **S3**) and sampled “open”, “semi-open” and “closed” conformations (**Figure S5**). Both experimental and computational studies suggested that flexibility of the ECL2 was important for activation and allosteric modulation of the A_1_AR.^39,44,45^ Therefore, highly flexibility of the ECL2 contributed to activation of the A_1_AR and receptor coupling to the G protein.

The A_2A_AR could couple to both the G_s_ and G_i_ proteins. This correlated with a recent experimental study that the A_2B_AR coupled with both the G_s_ and G_i_ proteins in human cells.^3^ The A_2B_AR was able to activate different downstream signaling pathways via different G proteins (including the G_s_ and G_i_) in the same cell type (e.g., HEK293 kidney and T24 bladder cancer cells) and couple to the same pathway via different G proteins in different cell types.^3^ Considering high similarity of A_2A_AR and A_2B_AR (72%), especially at the G protein coupling interface (**Figure S13**), we assume that the A_2A_AR would also couple to both the G_s_ and G_i_ proteins.

With low-energy conformations of AR-G protein complexes obtained from the GaMD simulations, further analysis revealed that complementary residue interactions were key for specific GPCR-G protein coupling. When coupling to different G proteins, one receptor could change its conformation and flexibility (notably in the TM6 intracellular domain), similarly for one G protein as coupled to different receptors (**Figures 1** and **6**). Provided highly complementary residue interactions at the interface, the A_2A_AR could strongly couple to the G_i_ protein in addition to the G_s_. However, coupling of the A_1_AR to the G_s_ protein became weaker than to the G_i_, due to significantly reduced residue interactions (**Figure 5**). The complementary residue interactions were identified to involve the receptor TM6, TM5, H8 and ICL2, as well as the Gα α5 helix, α4-β6 loop and αN-β1 loop. These regions have been highlighted to be important determinants for specific GPCR-G protein coupling in extensive experimental and computational studies as reviewed earlier.^10–12,46^

In summary, the GaMD simulations with unconstrained enhanced sampling and free energy calculations have provided important insights into the mechanism of specific G protein coupling to the A_1_AR and A_2A_AR. Nevertheless, effects of binding different extracellular ligands (e.g., agonists of varied potencies and allosteric modulators) on the GPCR-G protein interactions are subject to future studies. Furthermore, challenges remain to accurately predict the thermodynamic and kinetic properties of the G protein binding to the GPCRs in order to fully understand the dynamics of GPCR-G protein interactions. It is important to characterize both the association and dissociation pathways of the G protein coupling to GPCRs. Developments in computing power and enhanced simulation methodologies will be needed to address these problems in the future.

## Materials and Methods

### Gaussian accelerated molecular dynamics (GaMD)

GaMD enhances the conformational sampling of biomolecules by adding a harmonic boost potential to reduce the system energy barriers.^26^ When the system potential 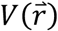 is lower than a reference energy E, the modified potential 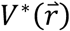 of the system is calculated as:

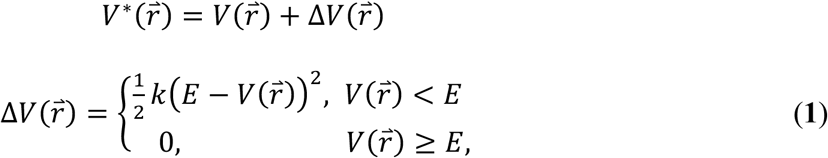

Where k is the harmonic force constant. The two adjustable parameters E and k are automatically determined on three enhanced sampling principles. First, for any two arbitrary potential values 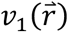 and 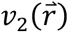 found on the original energy surface, if 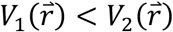, Δ*V* should be a monotonic function that does not change the relative order of the biased potential values; i.e., 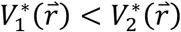. Second, if 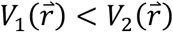, the potential difference observed on the smoothened energy surface should be smaller than that of the original; i.e., 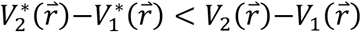. By combining the first two criteria and plugging in the formula of 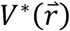 and Δ*V*, we obtain

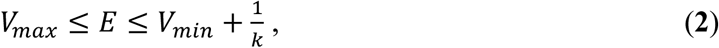

Where *V*_*min*_ and *V*_*max*_ are the system minimum and maximum potential energies. To ensure that **Eq. 2** is valid, *k* has to satisfy: *K* ≤ 1/(*V*_*max*_ − *V*_*min*_). Let us define: *K* = *K*_0_ ∙ 1/(*V*_*max*_ − *V*_*min*_), then 0 < *K*_0_ ≤ 1. Third, the standard deviation (SD) of Δ*V* needs to be small enough (i.e. narrow distribution) to ensure accurate reweighting using cumulant expansion to the second order: σ_ΔV_ =*K*(*E* − *V*_avg_)σ_D_ ≤ σ_0_, where *V*_avg_ and σ_v_ are the average and SD of Δ*V*with σ_0_ as a user-specified upper limit (e.g., 10*K*_B_*T*) for accurate reweighting. When E is set to the lower bound *E* = *V*_*max*_ according to **Eq. 2**, *K*_0_ can be calculated as

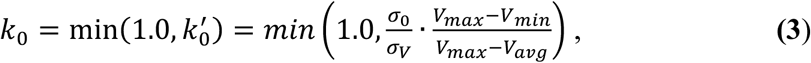

Alternatively, when the threshold energy E is set to its upper bound *E* = *V*_*min*_ + 1/*K*, *K*_0_is set to:

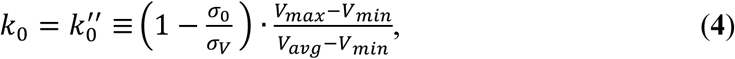

If 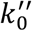 is calculated between 0 and 1. Otherwise, *K*_0_is calculated using **Eq. 3**.

### Energetic Reweighting of GaMD Simulations

For energetic reweighting of GaMD simulations to calculate potential of mean force (PMF), the probability distribution along a reaction coordinate is written as *P*∗(*A*). Given the boost potential Δ*V*(*r*) of each frame, *P*∗(*A*) can be reweighted to recover the canonical ensemble distribution *P*(*A*), as:

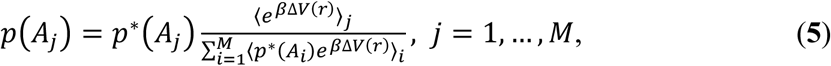

where *M* is the number of bins, *β* = *K*_B_*T* and 〈*e*^βΔV(r)〉^*j* is the ensemble-averaged Boltzmann factor of Δ*V*(*r*) for simulation frames found in the *j*^th^ bin. The ensemble-averaged reweighting factor can be approximated using cumulant expansion:

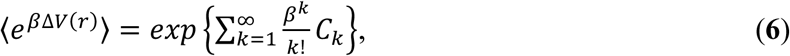

where the first two cumulants are given by:

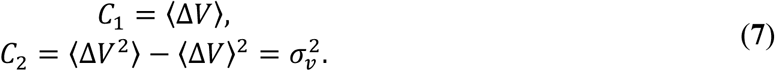

The boost potential obtained from GaMD simulations usually follows near-Gaussian distribution.^28^ Cumulant expansion to the second order thus provides a good approximation for computing the reweighting factor.^26,47^ The reweighted free energy *F*(*A*) = −*K*_B_*T* ln *P*(*A*) is calculated as:

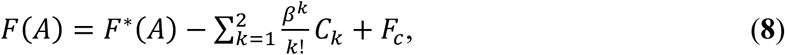

where *F*∗(*A*) = −*K*_B_*T* ln *P*∗(*A*) is the modified free energy obtained from GaMD simulation and *F*_c_ is a constant.

### System Setup

Cryo-EM structures of the ADO-A_1_AR-G_i_ (PDB: 6D9H)^8^ and NECA-A_2A_AR-G_s_ (PDB: 6GDG)^9^ were used for setting up simulation systems. Nanobody Nb35 in the cryo-EM structure of the NECA-A_2A_AR-G_s_ was deleted for simulation. In the 6GDG cryo-EM structure, residues that were missing in extracellular loop (ECL) 2 and the C terminus of the A_2A_AR were added using atomic coordinates obtained from the X-ray structure of the A_2A_AR bound by the mini-G_s_ protein (PDB: 5G53)^48^ after aligning the receptor transmembrane (TM) domain. Initial models of the ADO-A_1_AR-G_s_ and NECA-A_2A_AR-G_i_ protein complexes were obtained by switching the G proteins in the NECA-A_2A_AR-G_s_ and ADO-A_1_AR-G_i_ complexes after aligning the receptor TM domain (**Figure S1**). There was no clash between the ARs and G proteins.

According to previous findings, intracellular loop (ICL) 3 is highly flexible^21,49^ and removal of ICL3 does not appear to affect GPCR function. The ICL3 was thus omitted as in the cryo-EM structures for the simulations. In addition, helical domains of the G_i_ and G_s_ proteins missing in the cryo-EM structures were not included in the simulation models. This was based on earlier simulation of the β_2_AR-G_s_ complex, which showed that the helical domain fluctuated substantially.^21^ All chain termini were capped with neutral groups (acetyl and methylamide). All the disulphide bonds in the receptors and G proteins (i.e., Cys80^3.25^-Cys169^ECL2^ and Cys260^6.61^-Cys263^ECL3^ in the A_1_AR, Cys^743.22^-Cys146^ECL2^, Cys77^3.25^-Cys166^ECL2^, Cys71^ECL1^-Cys159^ECL2^ and Cys259^6.61^-Cys262^ECL3^ in the A_2A_AR, and Cys121-Cys149 in the G_β_ subunit of the G_s_ protein) that were resolved in the cryo-EM structures were maintained in the simulations. Using the *psfgen* plugin in VMD,^50^ protein residues were set to the standard CHARMM protonation states at neutral pH. For each of the complex systems, the receptor was inserted into a palmitoyl-oleoyl-phosphatidyl-choline (POPC) bilayer with all overlapping lipid molecules removed using the membrane plugin in VMD. The system charges were then neutralized at 0.15M NaCl using the *solvate* plugin in VMD.^50^ The simulation systems were summarized in **Table 1**, with an example computational model shown in **Figure S2**.

### Simulation Protocol

The CHARMM36 parameter set^51^ was used for the adenosine receptors, G proteins and POPC lipids. Force field parameters of agonists ADO and NECA were obtained from the CHARMM ParamChem web server.^52,53^ For each of the AR-G protein complex systems, initial energy minimization, thermalization, and 20ns cMD equilibration were performed using NAMD2.12^54^. A cutoff distance of 12 Å was used for the Van der Waals and short-range electrostatic interactions and the long-range electrostatic interactions were computed with the particle-mesh Ewald summation method. A 2-fs integration time step was used for all MD simulations and a multiple-time-stepping algorithm was used with bonded and short-range non-bonded interactions computed every time step and long-range electrostatic interactions every two time steps. The SHAKE algorithm was applied to all hydrogen-containing bonds. The NAMD simulation started with equilibration of the lipid tails. With all other atoms fixed, the lipid tails were energy minimized for 1,000 steps using the conjugate gradient algorithm and melted with a constant number, volume, and temperature (NVT) run for 0.5 ns at 310 K. The four systems were further equilibrated using a constant number, pressure, and temperature (NPT) run at 1 atm and 310 K for 10 ns with 5 kcal/(mol∙ Å^2^) harmonic position restraints applied to the protein and ligand atoms. The system volume was found to decrease with a flexible unit cell applied and level off with a 10-ns NPT run, suggesting that solvent and lipid molecules in the system were well equilibrated. Final equilibration of each system was performed using a NPT run at 1 atm pressure and 310 K for 0.5 ns with all atoms unrestrained. After energy minimization and system equilibration, conventional MD simulations were performed on each system for 20 ns at 1 atm pressure and 310 K with a constant ratio constraint applied on the lipid bilayer in the X-Y plane.

With the NAMD output structure, along with the system topology and CHARM36 force field files, the *ParmEd* tool in the AMBER package was used to convert the simulation files into the AMBER format.^55^ The GaMD module implemented in the GPU version of AMBER18^26,55^ was then applied to perform the GaMD simulation, which included a 8-ns short cMD simulation used to collect the potential statistics for calculating GaMD acceleration parameters, a 64-ns equilibration after adding the boost potential, and finally three independent 300-ns GaMD production simulations with randomized initial atomic velocities. All GaMD simulations were run at the “dual-boost” level by setting the reference energy to the lower bound. One boost potential is applied to the dihedral energetic term and the other to the total potential energetic term. The average and SD of the system potential energies were calculated every 800,000 steps (1.6 ns) for all simulation systems. The upper limit of the boost potential SD, σ0 was set to 6.0 kcal/mol for both the dihedral and the total potential energetic terms. Similar temperature and pressure parameters were used as in the NAMD simulations. A list of GaMD production simulations on the different ARs-G proteins complex systems is listed in **Table 1**.

### Simulation Analysis

CPPTRAJ^56^ and VMD^50^ were used to analyze the GaMD simulation trajectories. Particularly, distances were calculated between the Cα atoms of residues Arg^3.50^ and Glu^6.30^, the center-of-mass (COM) distance between the receptor NPxxY motif and the last 5 residues of the G_α_ α5 helix, and the COM distance between the G_α_ (excluding residues in the αN helix) and G_β_ (excluding residues 2-45 in the N-terminus) subunits. Root-mean-square fluctuations (RMSFs) were calculated for the protein residues and ligands, averaged over three independent GaMD simulations and color coded for schematic representation of each complex system (**Figure 1**). The representative low-energy conformations of AR-G protein were used to analyze the residue interaction contacts. The residue contact network between the AR and G protein was computed using van der Waals contacts between atoms, as described in Reference 37.^57^ For two-dimensional visualization, software Cytoscape^58^ was utilized to plot the residue contact network.

The PyReweighting^47^ toolkit toolkit was used to reweight distances, root-mean-square deviations (RMSDs) and the Gα α5 orientation angle to compute the potential of mean force (PMF) profiles. A bin size of 1.0 Å was used for the distances and RMSDs values, and 5.0 degree for the ^G_α_ α5 orientation angle. The cutoff was set to 500 frames for 2D PMF calculations. The 2D PMF^ profiles were obtained for each simulation system regarding agonist RMSD relative to the cryo-EM conformation and the AR:NPxxY-G:α5 distance (**Figure 2**), RMSD of the helix region in ECL2 relative to the cryo-EM structure and the AR:NPxxY-G:α5 distance (**Figures 4A and S5**), the distance between atom NE1 of W156 and atom O of G163 and RMSD of the helix region in ECL2 relative to the cryo-EM structure (**Figure S6**), the Arg^3.50^-Glu^6.30^ and the AR:NPxxY-G:α5 distances (**Figures 4B and S7**), the G_α_ α5 orientation angle and the Arg^3.50^-Glu^6.30^ distance (**Figures 4C and S9**), and increase of the G_α_-G_β_ distance and the Arg^3.50^-Glu^6.30^ distance (**Figures 4D and S10**). Time courses of these reaction coordinates obtained from the GaMD simulation were plotted in **Figures S14-S17**.

## ASSOCIATED CONTENT

## AUTHOR INFORMATION

### Author Contributions

Y.M. designed research; J.W. performed research; J.W. and Y.M. analyzed data; and J.W. and Y.M. wrote the paper.

### Notes

The authors declare no competing financial interest.

## Acknowledgements

We thank Waseem Ahmad for proofreading the manuscript. Computing time was provided on the Comet and Stanford EXtream supercomputers through the Extreme Science and Engineering Discovery Environment award TG-MCB180049 and the Edison and Cori supercomputers through the National Energy Research Scientific Computing Center project M2874. This work was supported in part by the American Heart Association (Award 17SDG33370094) and the startup funding in the College of Liberal Arts and Sciences at the University of Kansas.

