## Supporting Information for "Mechanistic Insights into Specific G Protein Interactions with Adenosine Receptors Revealed by Accelerated Molecular Simulations"

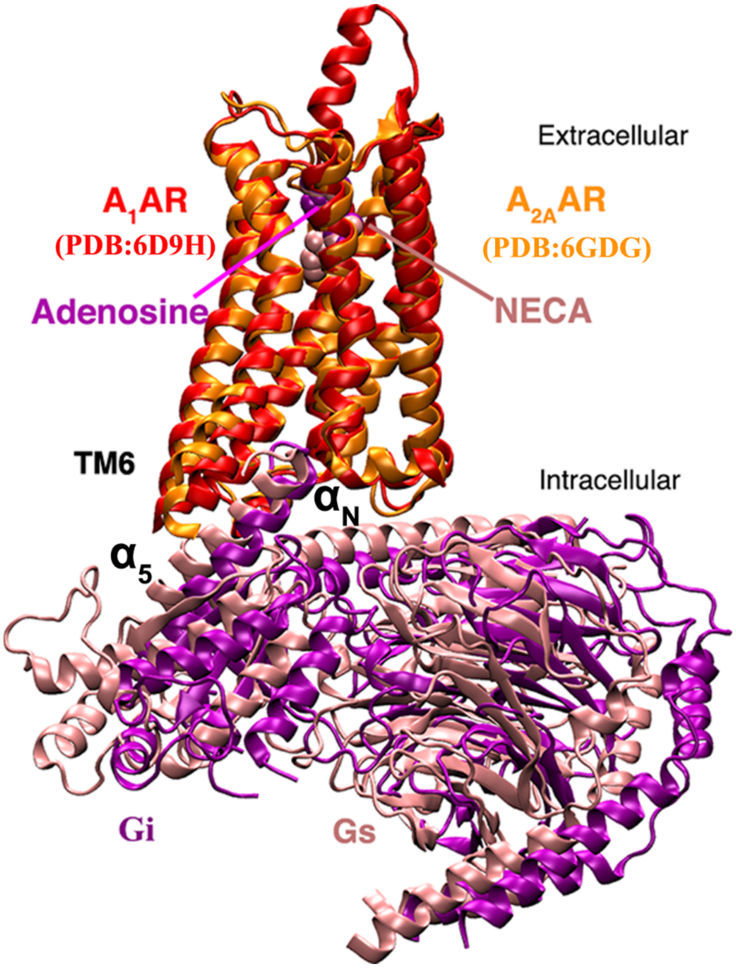


**Figure S1**. Comparison of cryo-EM structures between the $A_1AR$-$G_i$ (PDB: 6D9H) and $A_{2A}AR$-$G_s$ protein (PDB: 6GDG) complexes.

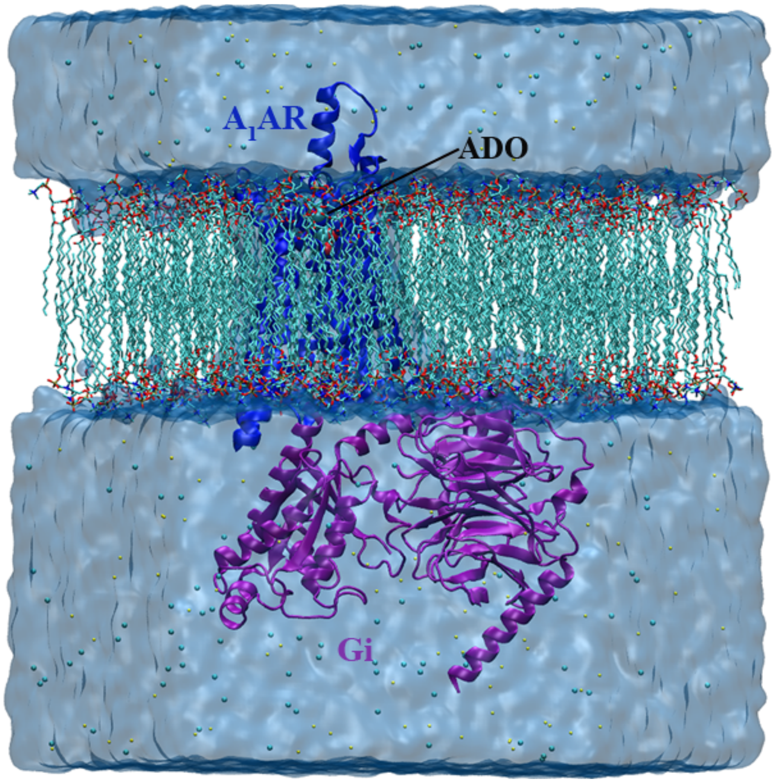


**Figure S2**. Schematic representation of the computational model of adenosine receptor-G protein complex systems as shown for the ADO-$A_1$AR-$G_i$. The receptor was inserted into a POPC bilayer and solvated in an aqueous medium of 0.15 M NaCl. The ADO agonist atoms are shown in spheres and colored by atom names. The receptor and G protein are shown in blue and purple, respectively.

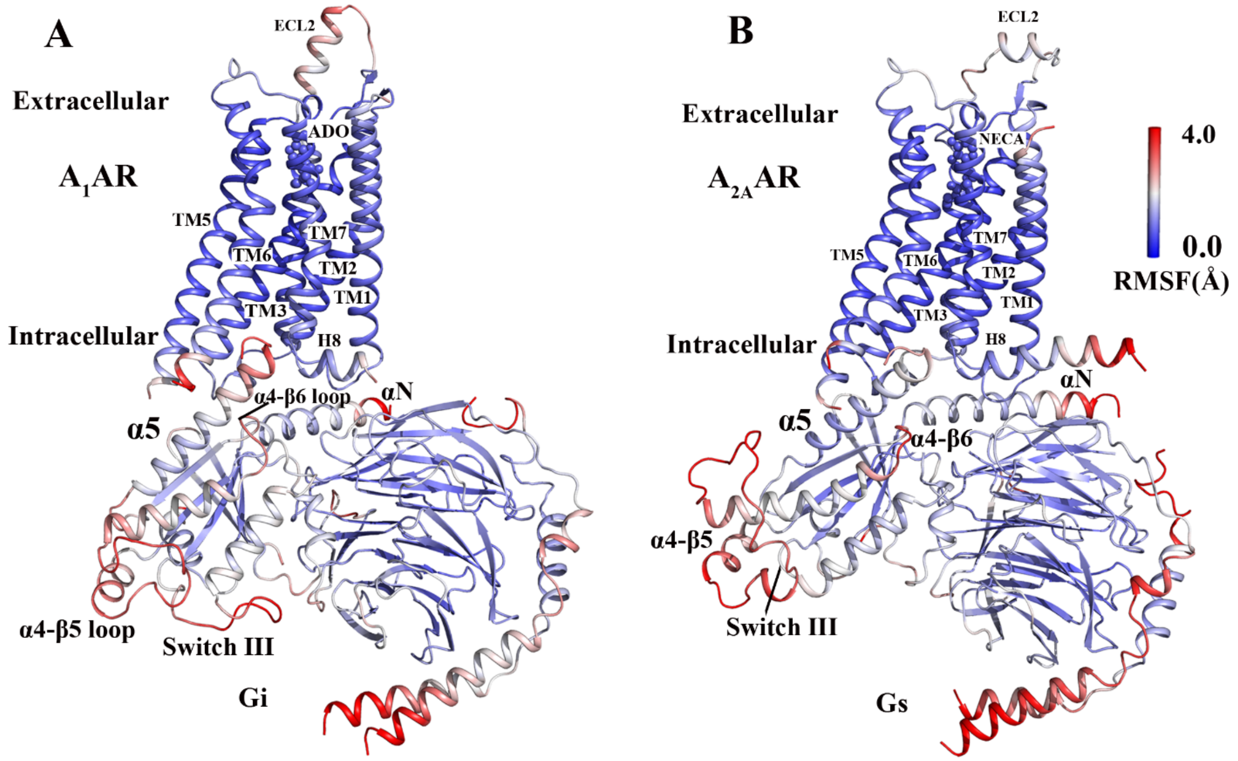


**Figure S3**. Comparison of structural flexibility of active adenosine receptor-G protein complexes obtained from GaMD simulations: **(A)** the ADO-$A_1$AR-$G_i$ protein and **(B)** NECA-$A_2$AR-$G_s$ protein complex systems, which are colored by root-mean square fluctuations (RMSFs). A color scale of 0 Å (blue) to 4 Å (red) is used.

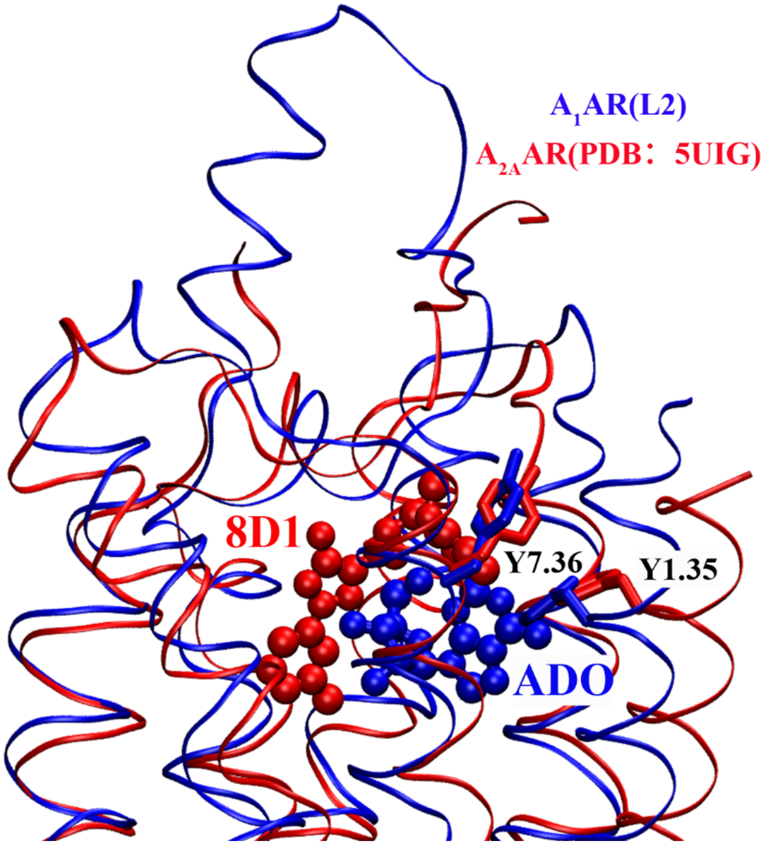


**Figure S4**. The “L2” representative conformation of ADO in the ADO-$A_1$AR-$G_s$ complex (blue) in compared with the X-ray structure of the $A_{2A}$AR bound by antagonist 8D1 (PDB: 5UIG, red).

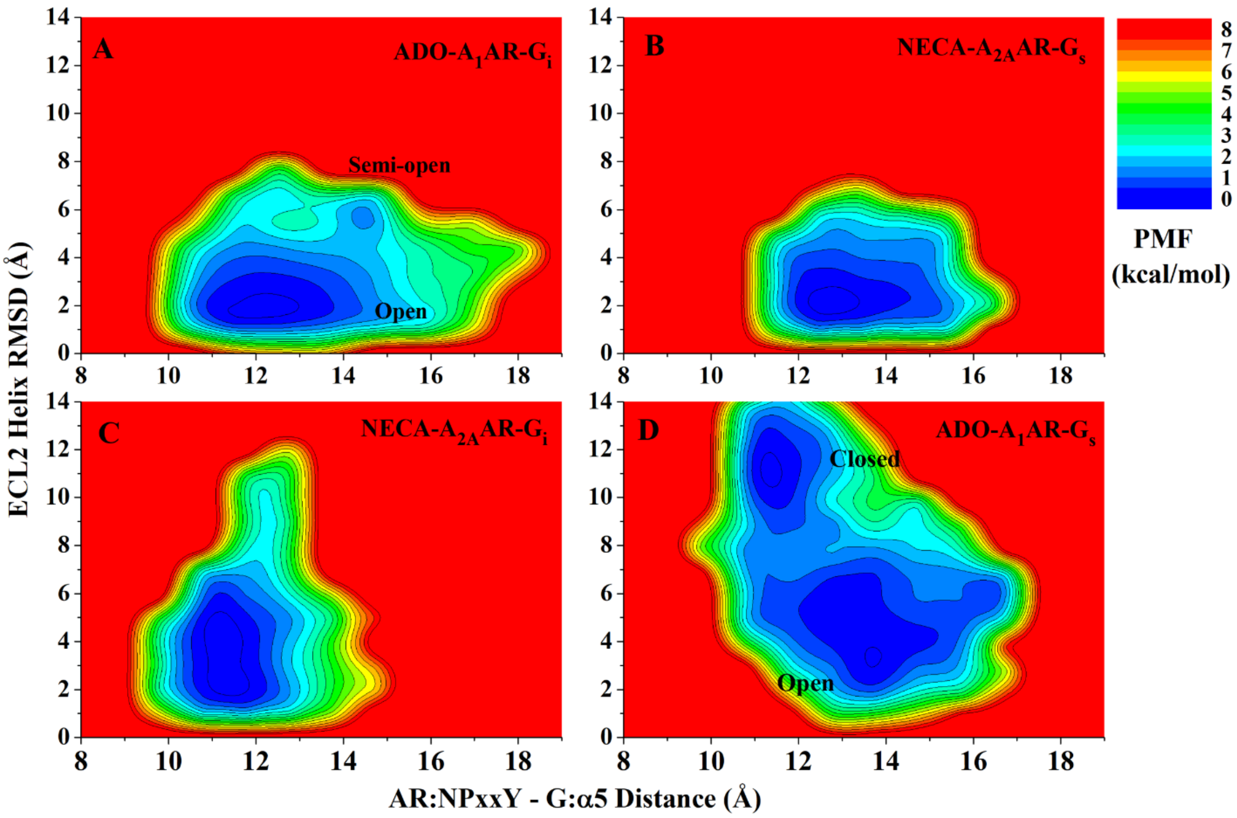


**Figure S5.** Distinct low-energy conformational states of the ECL2 were sampled by the adenosine receptor-G protein complexes. 2D PMF profiles of the **(A)** ADO-$A_1$AR-$G_i$, **(B)** NECA-$A_{2A}$AR-$G_s$, **(C)** NECA-$A_{2A}$AR-$G_i$ and **(D)** ADO-$A_1$AR-$G_s$ complex systems regarding RMSD of the helix region in ECL2 relative to the cryo-EM structure and the distance between COMs of the receptor NPxxY motif and the last 5 residues of the Gα α5 helix.

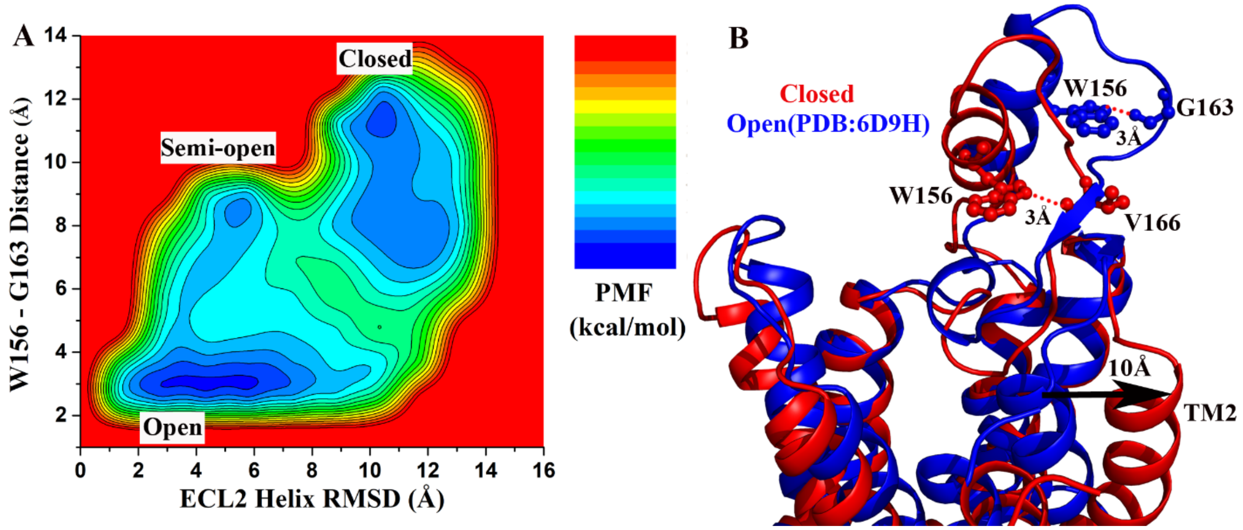


**Figure S6**. **(A)** 2D PMF profiles of the ADO-$A_1$AR-$G_s$ complex system regarding RMSD of the helix region in ECL2 relative to the cryo-EM structure and the distance between the atom NE1 of W156 and atom O of G163 in the $A_1$AR ECL2. **(B)** The representative conformations of open (cryo-EM structure 6D9H, blue) and closed (red) states of ECL2 in the ADO-$A_1$AR-$G_s$ protein complex.

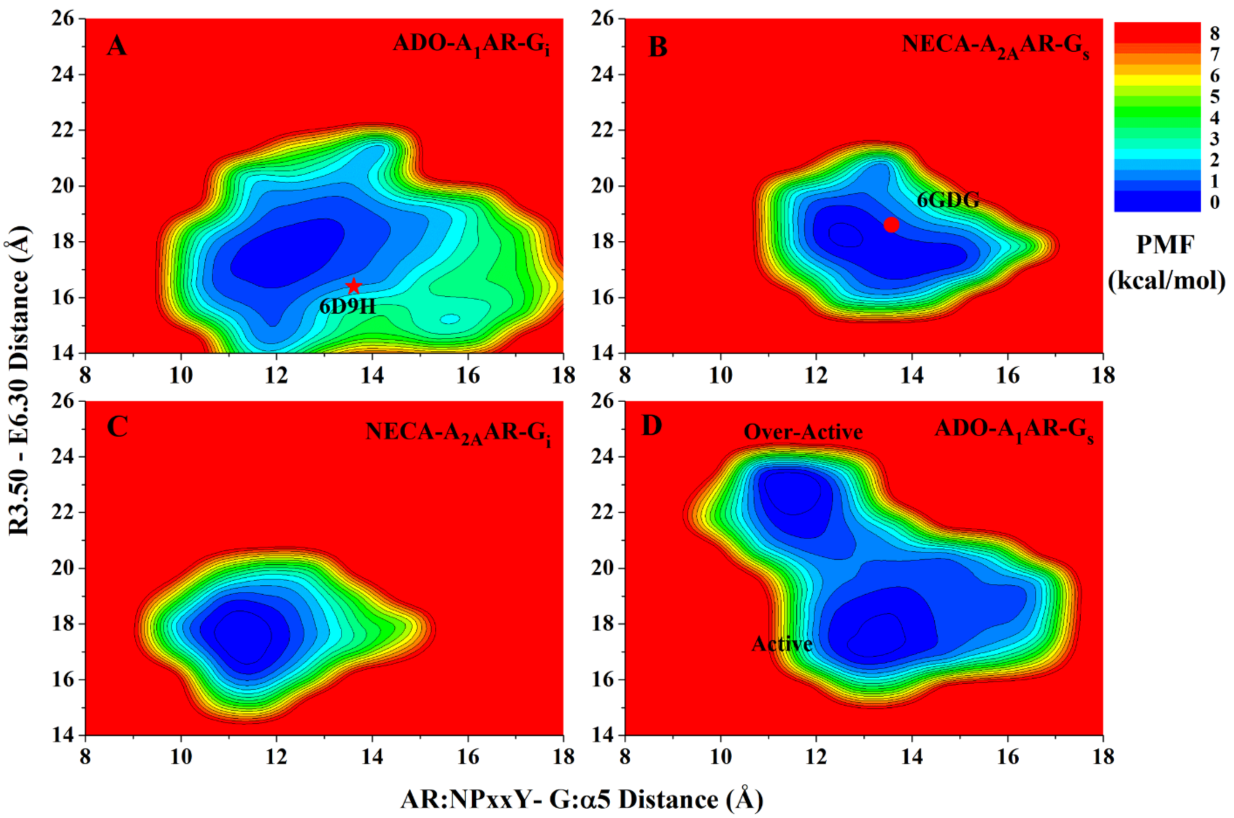


**Figure S7**. 2D PMF profiles of the **(A)** ADO-$A_1$AR-$G_i$, **(B)** NECA-$A_{2A}$AR-$G_s$, **(C)** NECA-$A_{2A}$AR-$G_i$ and **(D)** ADO-$A_1$AR-$G_s$ complex systems regarding the distance between the Cα atoms of residues $Arg^{3.50}$ and $Glu^{6.30}$ in the receptors and the distance between COMs of the receptor NPxxY motif and the last 5 residues of the Gα α5 helix. The red star and dot indicate the ADO-$A_1$AR-$G_i$ (6D9H) and NECA-$A_{2A}$AR-$G_s$ (6GDG) cryo-EM structures, respectively.

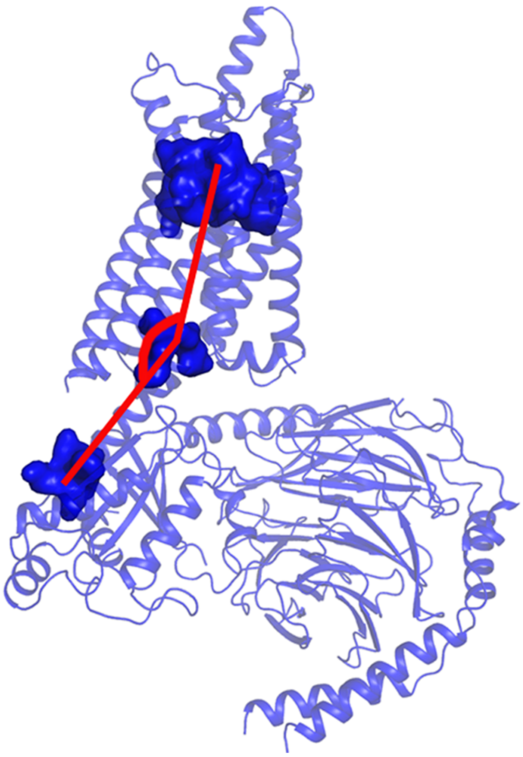

**Figure S8**. Illustration of the orientation angle of the Gα α5 helix, which is defined as the angle between COMs of the receptor orthosteric pocket, the last 5 and first 5 residues of the Gα α5 helix.

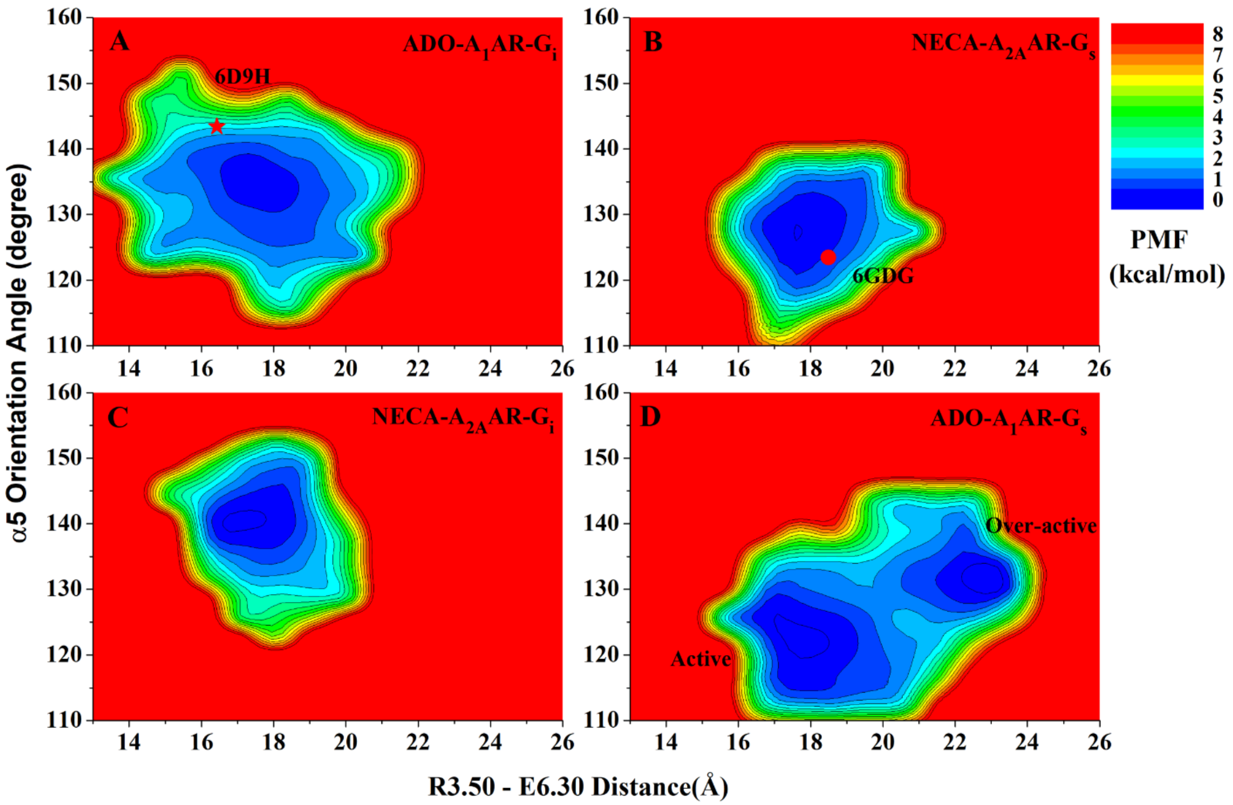


**Figure S9**. 2D PMF profiles of the **(A)** ADO-$A_1$AR-$G_i$, **(B)** NECA-$A_{2A}$AR-$G_s$, **(C)** NECA-$A_{2A}$AR-$G_i$ and **(D)** ADO-$A_1$AR-$G_s$ complex systems regarding orientation angle of the Gα α5 helix (illustrated in **Fig. S8**) and the distance between the Cα atoms of residues $Arg^{3.50}$ and $Glu^{6.30}$ in the receptors. The red star and dot indicate the ADO-$A_1$AR-$G_i$ (6D9H) and NECA-$A_{2A}$AR-$G_s$ (6GDG) cryo-EM structures, respectively.

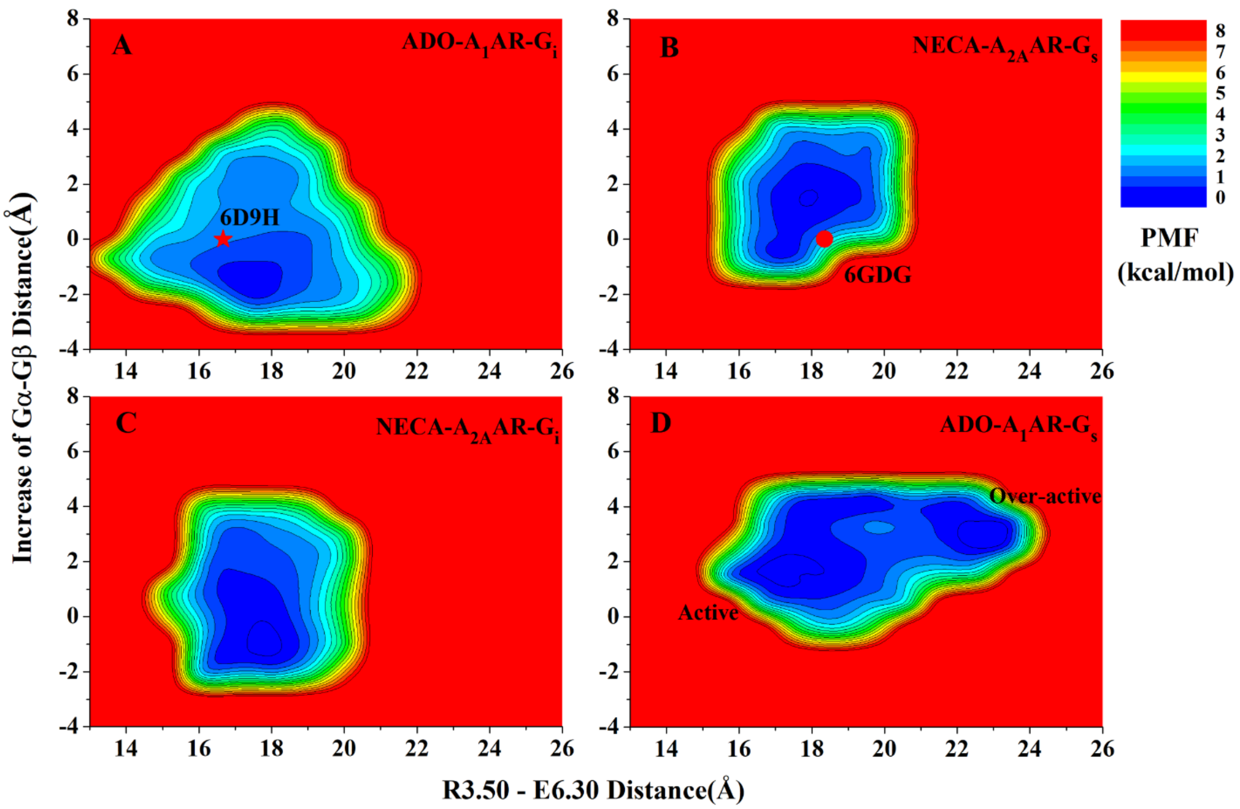


**Figure S10**. Distinct low-energy conformational states of the G proteins observed in GaMD simulations: 2D PMF profiles regarding the increase in the distance between the COMs of $G_{\alpha}$ and $G_{\beta}$ subunits and the distance between the Cα atoms of residues $Arg^{3.50}$ and $Glu^{6.30}$ in the receptors in the **(A)** ADO-$A_1$AR-$G_i$, **(B)** NECA-$A_{2A}$AR-$G_s$, **(C)** NECA-$A_{2A}$AR-$G_i$ and **(D)** ADO-$A_1$AR-$G_s$ complex systems, respectively. The red star and dot indicate cryo-EM structures of the ADO-$A_1$AR-$G_i$ (6D9H) and NECA-$A_{2A}$AR-$G_s$ (6GDG), respectively. In comparison with the cryo-EM structures, the $G_{\alpha}$ and $G_{\beta}$ subunits in the $G_i$ protein were induced to move closer, while those in the $G_s$ protein tended dissociate from each other.

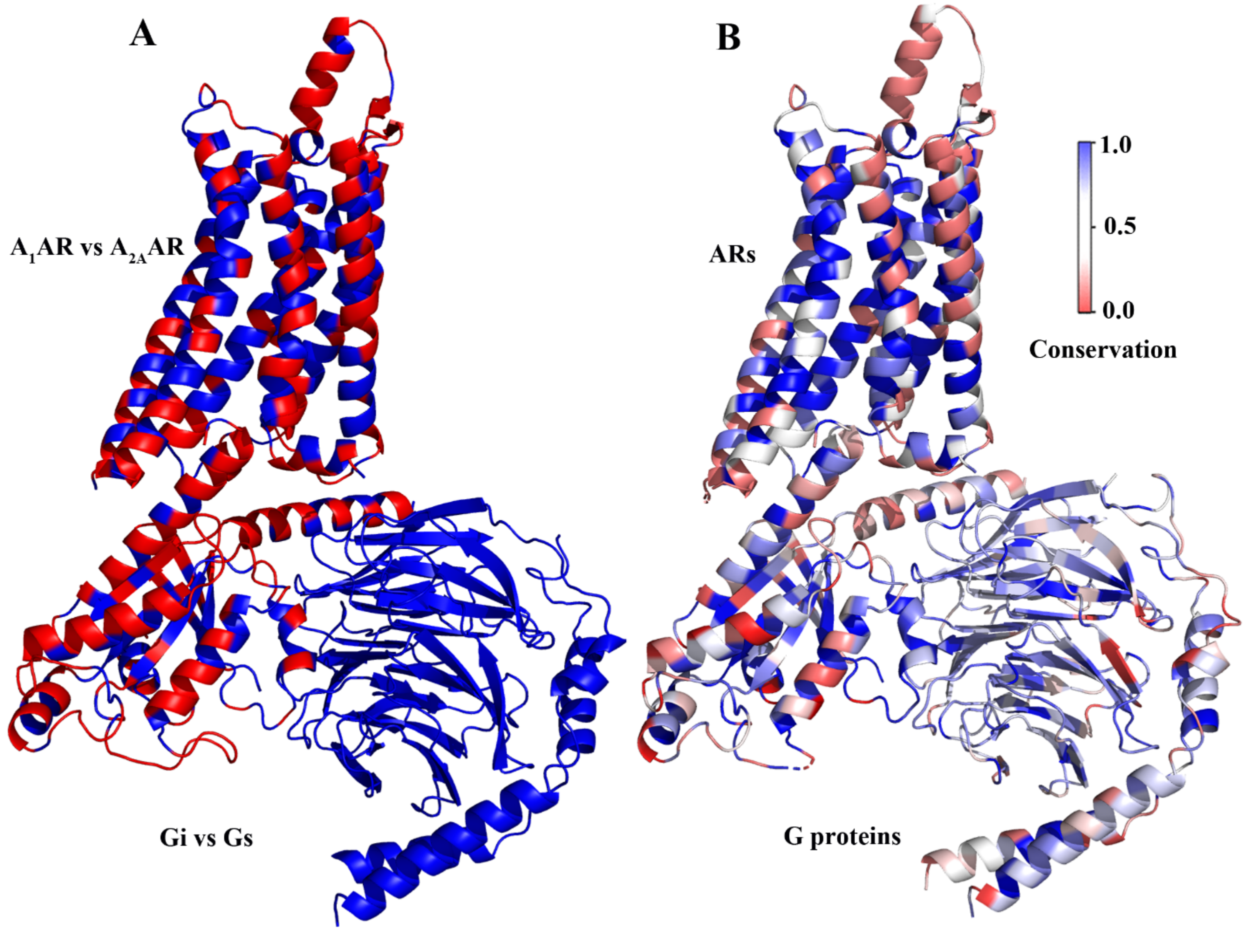


**Figure S11**. (**A**) Schematic representation of the $A_1AR$ colored by sequence identity among the $A_1AR$ and $A_{2A}AR$. (**B**) Schematic representation of the $A_1AR$-Gi complex colored by sequence conservation across four subtypes of ARs, and 16, 7, and 12 subtypes of the $G_\alpha$, $G_\beta$ and $G_\gamma$ subunits, respectively.

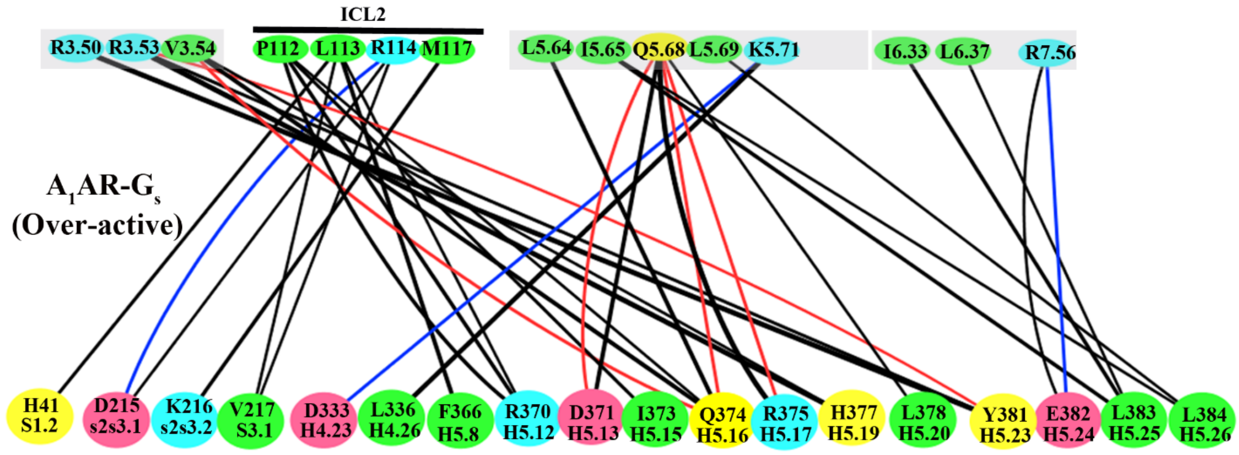


**Figure S12.** Residue interactions between the $G_\alpha$ and receptor in the "Over-active" state of the ADO-$A_1$AR-$G_i$.

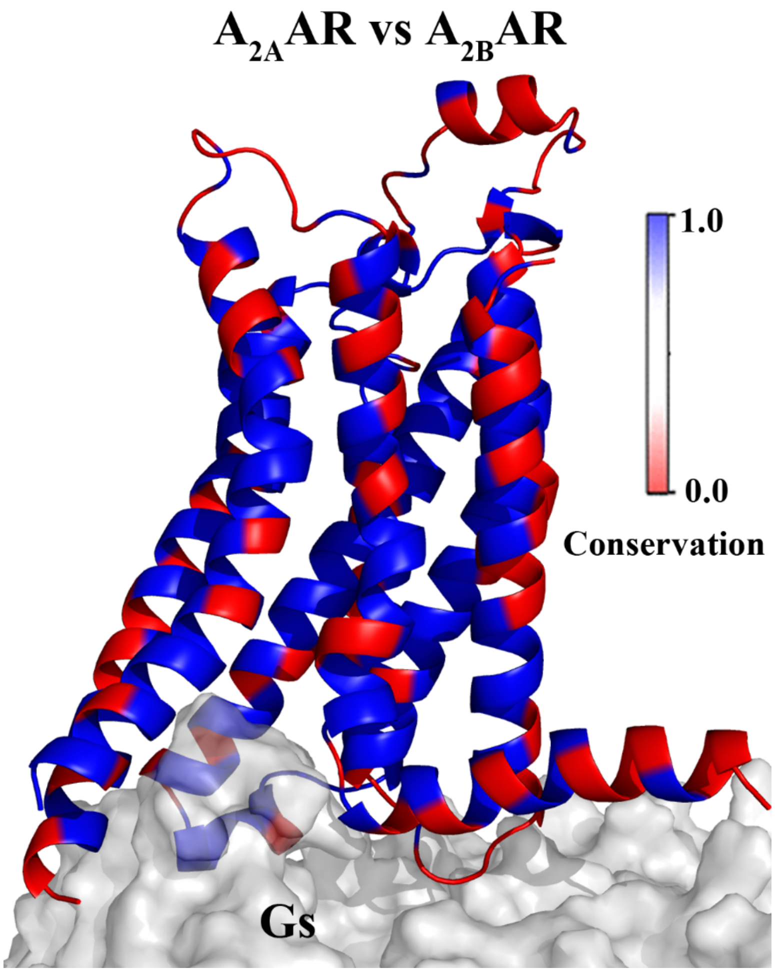


**Figure S13**. Schematic representation of the $A_{2A}$AR colored by sequence conservation between the $A_{2A}$AR and $A_{2B}$AR.

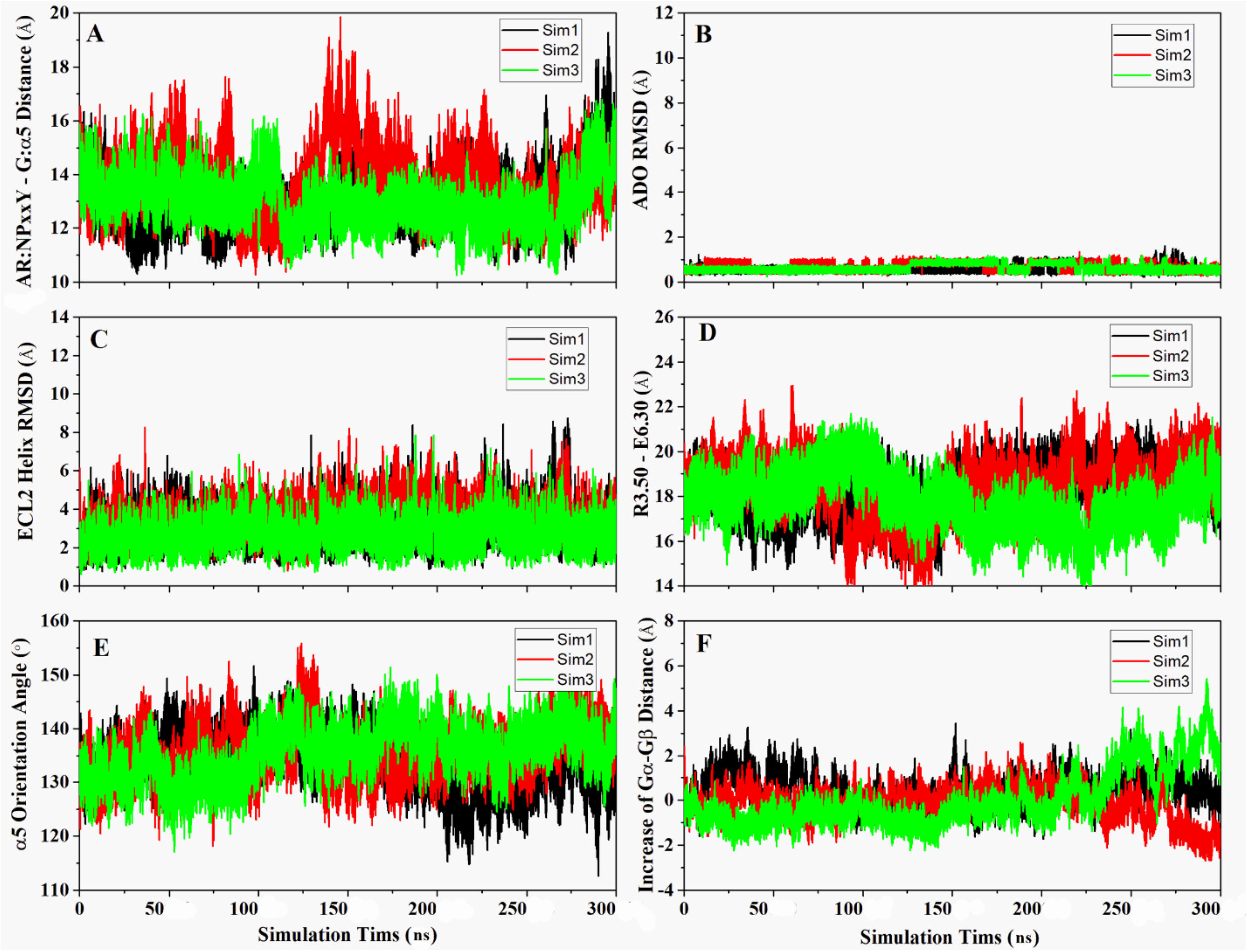


**Figure S14**. GaMD simulations of the ADO-$A_1$AR-$G_i$ system: Time courses of **(A)** $A_1$AR:NPxxY-G:α5 distance, **(B)** agonist RMSD, **(C)** ECL2 helix RMSD, **(D)** R3.50-E6.30 distance, **(E)** α5 orientation angle, and **(F)** increase of $G_\alpha$-$G_\beta$ distance.

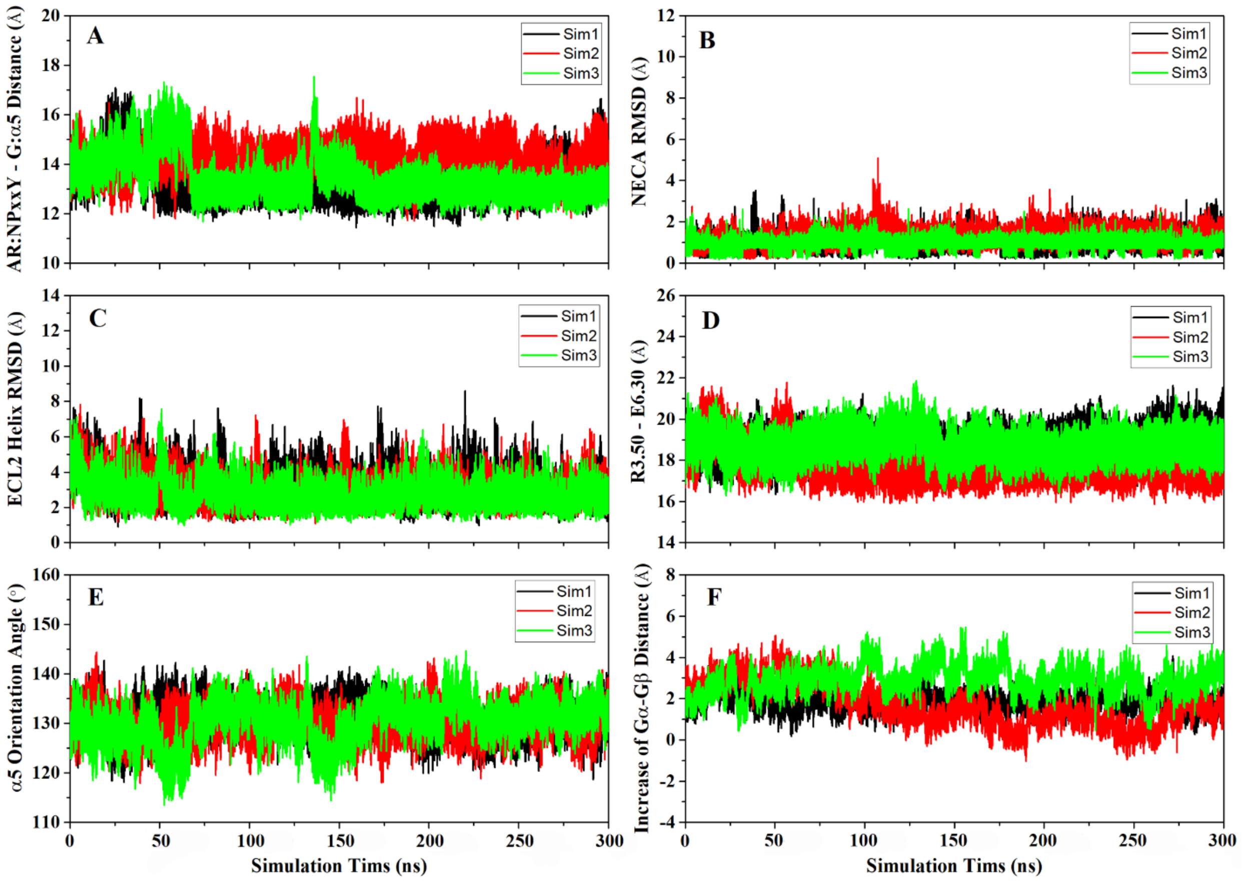


**Figure S15**. GaMD simulations of the NECA-$A_{2A}$AR-$G_s$ system: time courses of **(A)** $A_{2A}$AR:NPxxY-G:α5 distance, **(B)** agonist RMSD, **(C)** ECL2 helix RMSD, **(D)** R3.50-E6.30 distance, **(E)** α5 orientation angle, and **(F)** increase of $G_\alpha$-$G_\beta$ distance.

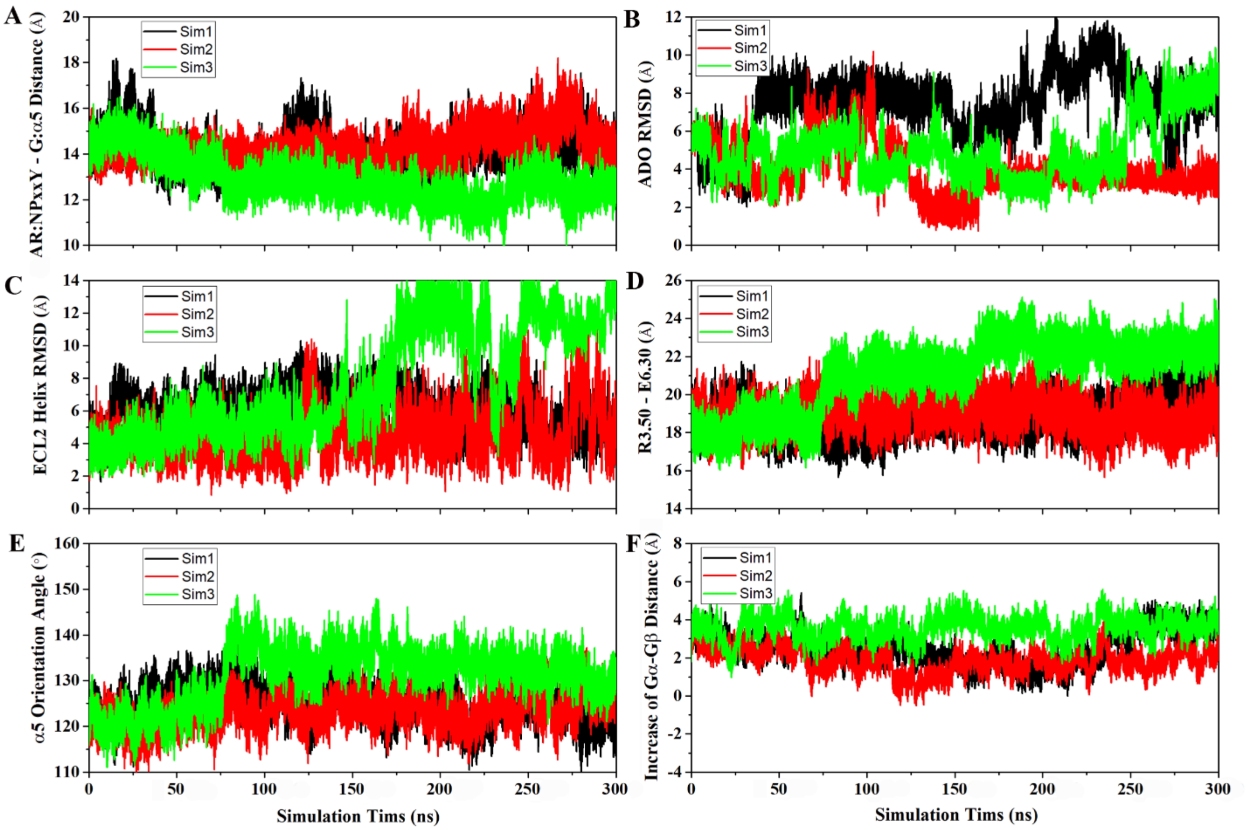


**Figure S16**. GaMD simulations of the ADO-$A_1$AR-$G_s$ system: time courses of **(A)** $A_1$AR:NPxxY-G:α5 distance, **(B)** agonist RMSD, **(C)** ECL2 helix RMSD, **(D)** R3.50-E6.30 distance, **(E)** α5 orientation angle, and **(F)** increase of $G_\alpha$-$G_\beta$ distance.

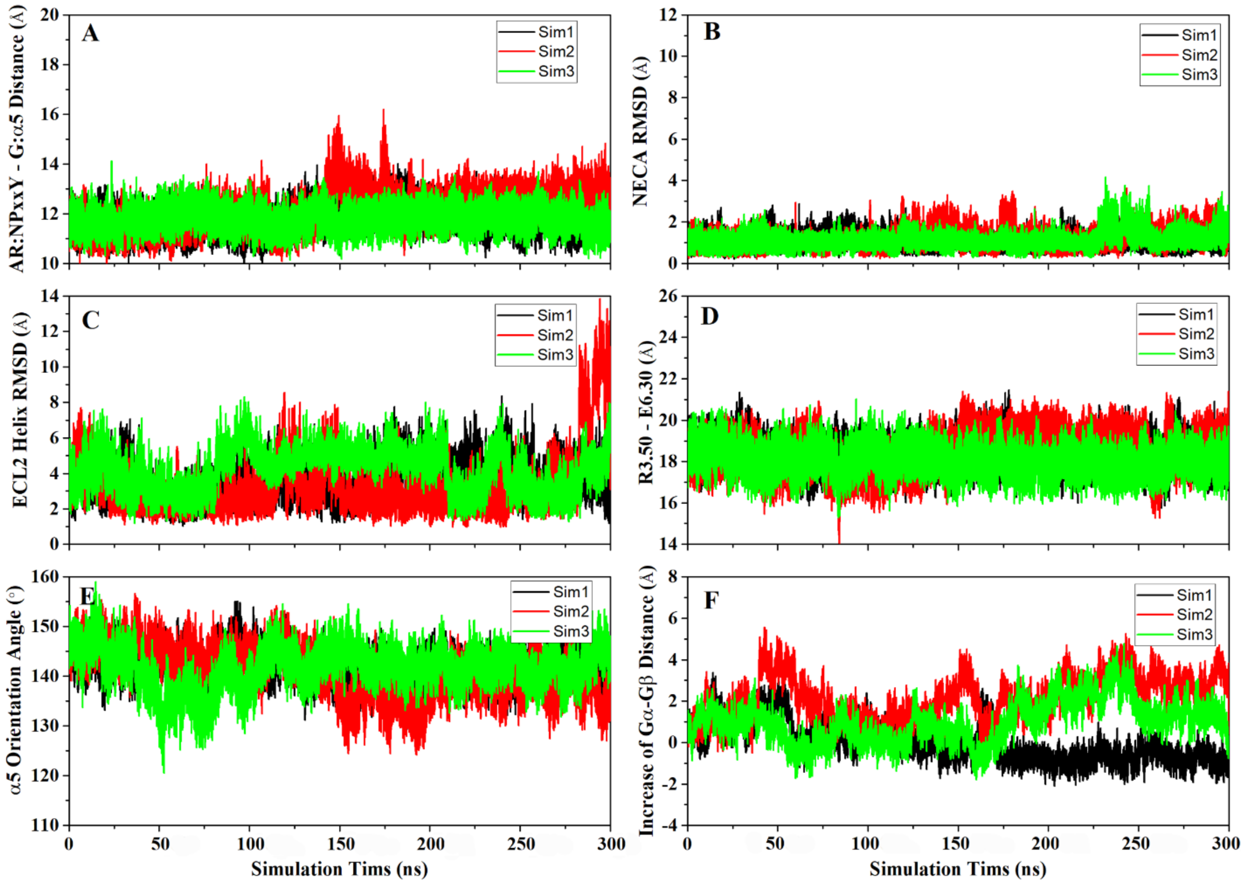


**Figure S17**. GaMD simulations of the NECA-$A_{2A}$AR-$G_i$ system: time courses of **(A)** $A_{2A}$AR:NPxxY-G:α5 distance, **(B)** agonist RMSD, **(C)** ECL2 helix RMSD, **(D)** R3.50-E6.30 distance, **(E)** α5 orientation angle, and **(F)** increase of $G_\alpha$-$G_\beta$ distance.
